# Insight in the quorum sensing-driven lifestyle of the non-pathogenic *Agrobacterium tumefaciens* 6N2 and the interactions with the yeast *Meyerozyma guilliermondii*

**DOI:** 10.1101/2021.10.17.464673

**Authors:** Elisa Violeta Bertini, Mariela Analía Torres, Thibaut Léger, Camille Garcia, Kar-Wai Hong, Teik Min Chong, Lucía I. Castellanos de Figueroa, Kok-Gan Chan, Yves Dessaux, Jean-Michel Camadro, Carlos Gabriel Nieto-Peñalver

**Affiliations:** PROIMI, CONICET (Planta Piloto de Procesos Industriales Microbiológicos), Av. Belgrano y Pje. Caseros, Tucumán, Argentina; ProteoSeine@IJM, Université de Paris, CNRS, Institut Jacques Monod, F-75006 Paris, France; Institute of Biological Sciences, Faculty of Science, University of Malaya, 50603 Kuala Lumpur, Malaysia; International Genome Centre, Jiangsu University, 212013, Zhenjiang, China; Institute of Marine Sciences, Shantou University, Shantou 515063, China; Instituto de Microbiología, Facultad de Bioquímica, Química y Farmacia, Universidad Nacional de Tucumán, Tucumán, Argentina; Department of Biotechnology, Faculty of Applied Sciences, UCSI University, Cheras, Wilayah Persekutuan Kuala Lumpur, Malaysia; Institut de Biologie Intégrative de la Cellule, Université Paris-Sud, Université Paris-Saclay, CNRS, CEA,, F-91190 Gif-sur-Yvette, France; CNRS, Institut Jacques Monod, Univ. Paris Diderot, Paris, France

**Keywords:** QUORUM SENSING, INTERACTIONS, ENDOPHYTIC, PROTEOMIC, AGROBACTERIA

## Abstract

*Agrobacterium tumefaciens* is considered a prominent phytopathogen, though most isolates are nonpathogenic. Agrobacteria can inhabit plant tissues interacting with other microorganisms. Yeasts are likewise part of these communities. We analyzed the quorum sensing (QS) systems of *A. tumefaciens* strain 6N2, and its relevance for the interaction with the yeast *Meyerozyma guilliermondii*, both sugarcane endophytes. We show that strain 6N2 is nonpathogenic, produces OHC8-HSL, OHC10-HSL, OC12-HSL and OHC12-HSL as QS signals, and possesses a complex QS architecture, with one truncated, two complete systems, and three additional QS-signal receptors. A proteomic approach showed differences in QS-regulated proteins between pure (64 proteins) and dual (33 proteins) cultures. Seven proteins were consistently regulated by quorum sensing in pure and dual cultures. *M. guilliermondii* proteins influenced by QS activity were also evaluated. Several up- and down- regulated proteins differed depending on the bacterial QS. These results show the importance of the QS regulation in the bacteria-yeast interactions.

**Highlights:** The avirulent *A. tumefaciens* 6N2 has two replicons and a complex QS architecture

The profile of QS-regulated proteins is modified in dual cultures with *Pa. laurentii*

The bacterial QS activity alters the proteome of the yeast *Pa. laurentii*

## Introduction

*Agrobacterium tumefaciens* is an alpha-proteobacterium of the Rhizobiaceae family, considered as one of the most important plant pathogens, which produces characteristic crown galls on numerous dicotyledoneous plants [1]. Its pathogenicity is related to the transfer of a piece of DNA, the T-DNA, from its oncogenic Ti plasmid, to the plant cell. However, in nature, most agrobacterial strains are devoid of a Ti plasmid, and are in consequence avirulent commensals [2]. The conjugation of Ti plasmid depends partially on a quorum sensing (QS) -regulated process [3]. QS is a cell–cell communication system that coalesces gene expression with the bacterial cell concentration [4]. It relies upon the production by LuxI homolog enzymes of signal molecules, termed autoinducers, whose concentration theoretically mimics that of the producing bacteria [5]. QS signals are perceived by a complementary LuxR homolog receptor protein when signals, hence cells, reach a threshold concentration [5]. Once the sensor binds the signal, it becomes activated and modifies the expression of QS-target genes. The model *A. fabrum* (formerly *A. tumefaciens*) strain C58 possesses a LuxI/LuxR-type QS system that utilizes 3-oxo-*N*- octanoyl-homoserine lactone (3OC8-HSL) as QS signal [6]. 3OC8-HSL, a member of the acyl homoserine lactone (AHL) family, the most characterized QS molecules in proteobacteria, is synthesized by the TraI enzyme, and is bound by the TraR receptor. The 3OC8-HSL-TraR complex activates the transcription of genes involved in the conjugative transfer of the Ti plasmid [7]. Although largely characterized in the strain C58 and other pathogenic strains, little is known about QS systems in commensal agrobacteria.

Though mostly considered a soil inhabitant, it is now clear that agrobacteria can also colonize the inner plant tissues, living as endophytes in stems, fruits and roots [8,9]. To date, their interactions with the host and other microorganisms in those particular niches remains poorly evaluated. Noteworthy, yeasts are also part of these complexes communities. Ascomycetous and Basidiomycetous yeasts have been identified as endophytes, including *Candida, Rhodotorula, Cryptococcus, Hanseniaspora, Debaryomyces* and *Metschnikowia*. It is expectable that these unicellular fungi interact with bacteria, including agrobacteria, in the endophytic polymicrobial communities. Their role in QS mediated interactions is unknown, even if a capacity to inactivate AHLs was demonstrated in several species [10].

During a previous survey of the endophytic microbiota of sugarcane (*Saccharum officinarum* L.), we isolated the yeast *Meyerozyma guilliermondii* strain 6N and *A. tumefaciens* strain 6N2 from the same node section, suggesting that these two microorganisms can co-occupy this niche and, in consequence, interact with each other [11,12]. In contrast to other species, this *M. guilliermondii* isolate show a very weak capacity to inactivate AHLs [10].

Information on the influence of the QS regulatory mechanisms on the interkingdom interactions remains scarce. Especially, little is known about how the QS regulation of a microorganism can affect the physiology of a second microorganism. In this report, we describe the complex architecture of the *A. tumefaciens* 6N2 QS system, responsible for the production of several AHLs. We performed proteomic analyses to characterize the QS regulation in this strain, and unveil how it is influenced in a dual culture with *M. guilliermondii* 6N and how this second microorganism is affected by the bacterial QS activity.

## Results

### Strain 6N2 is a bona fide *Agrobacterium tumefaciens* isolate producing several AHLs

Strain 6N2 showed a 16S rDNA sequence highly similar to those of the *Agrobacterium*/*Rhizobium* group (Genbank accession number MG062741). The sugarcane plant utilized in its isolation presented no symptoms of tumor formation, suggesting the non-pathogenicity of this isolate. This was confirmed with *A. thaliana* and tomato plants, which did not develop the characteristic tumors after inoculation with 6N2 strain (Fig. 1). The fragmentation of molecules obtained from culture extracts confirmed the production of AHLs by strain 6N2, according to the characteristic [M+H]^+^ of 102 m/z (Fig. 2). The determination of parent ions showed 4 molecules of [M+H]^+^ 244.4, 272.5, 298.6 and 300.6 m/z (Fig. 2), attributed to *N*-3-hydroxy-octanoyl-homoserine lactone (OHC8-HSL), *N*-3-hydroxy-decanoyl-homoserine lactone (OHC10-HSL), *N*-3-oxo-dodecanoyl-homoserine lactone (OC12-HSL) and *N*-3-hydroxy-dodecanoyl-homoserine lactone (OHC12-HSL), respectively (Suppl. Fig. 1).

**Figure 1.**
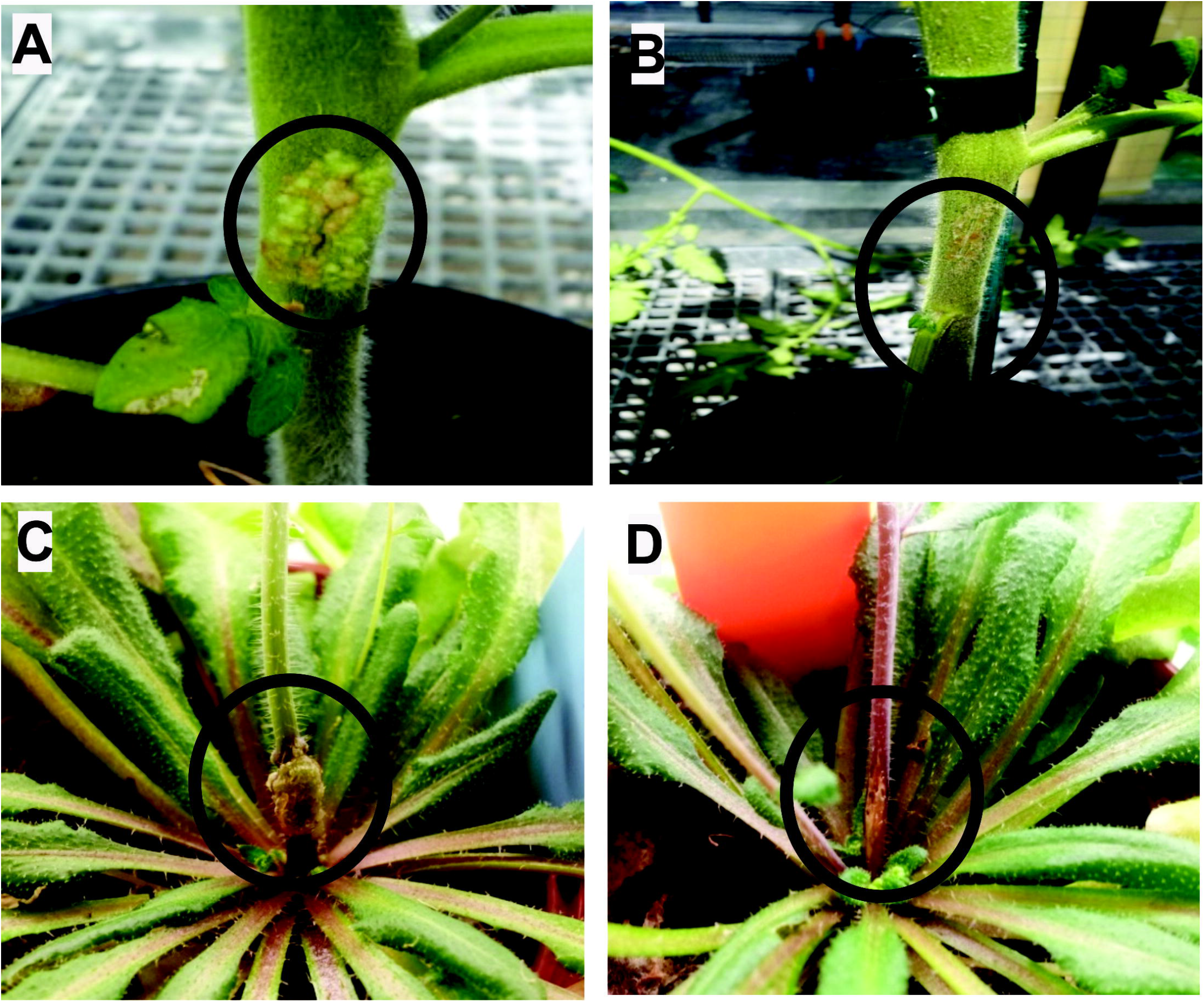
Pathogenesis tests on model plants. *A. tumefaciens* 6N2 did not develop the characteristic tumors on tomato (B) and *A. thaliana* (D) plants. A and C show the corresponding controls with *A. fabrum* C58.

**Figure 2.**
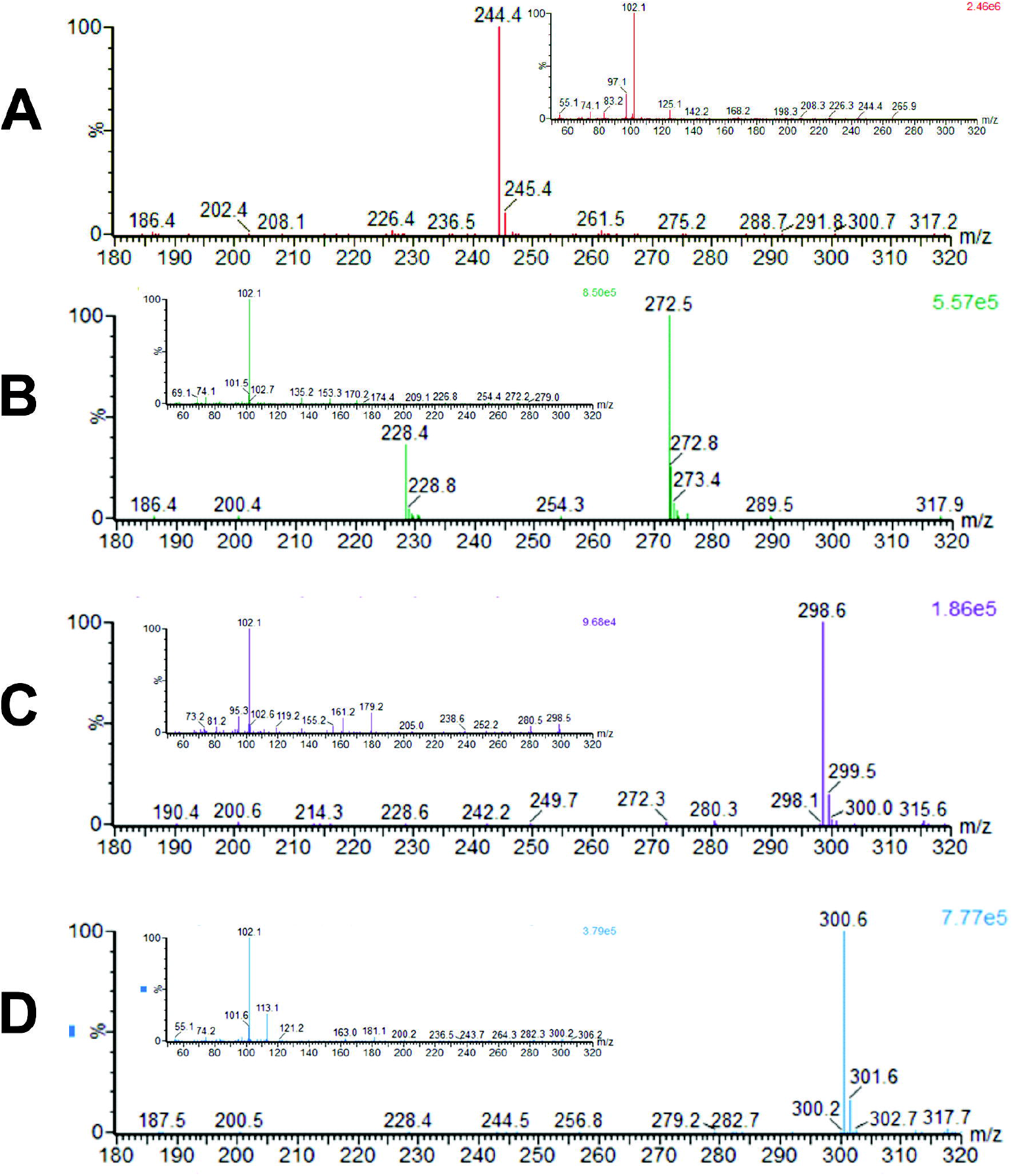
Mass spectrometric identification of AHLs produced by *A. tumefaciens* 6N2. The analysis of supernatant extracts showed the presence of molecules of [M+H]^+^ 244.4 (A), 272.5 (B), 298.6 (C) and 300.6 (D) m/z compatible with OHC8-HSL, OHC10-HSL, OC12-HSL and OHC12-HSL, respectively. In all the cases, the fragmentation produced a characteristic [M+H]^+^ of 102 m/z. See structures in Suppl. Fig. 1.

### Genomic characterization of *A. tumefaciens* 6N2

Genome sequencing of strain 6N2 revealed 2 replicons of 2,913,790 bp and 2,168,919 bp (Fig. 3A and B). The second replicon was assumed to be a linear chromosome considering the cumulative GC skew that suggested a replication origin at the center of the sequence (data not shown), and the identification of a *telA* ortholog (AT6N2_L1435), coding for TelA protelomerase. Genome annotation produced 3,013 and 2,074 CDS in the circular and the linear chromosome, respectively (Fig. 3). No traces of Ti or At plasmids were detected. Prophage *16-3* genes (coordinates: 282,851-343,277) were detected in the circular chromosome; several incomplete prophages (RHEph01, RcCronus, XcP1, SH2026Stx1 and Stx2a_F451) were predicted in the circular and linear chromosome (data not shown). Type IV (T4SS) and VI (T6SS) secretion systems were identified in the linear chromosome (Fig. 3B). Genomic islands were predicted in both chromosomes (Fig. 3A and B), and a probable integrative and conjugative element (ICE) in the linear chromosome (coordinates: 712,734-940,892) (Fig. 3B).

**Figure 3.**
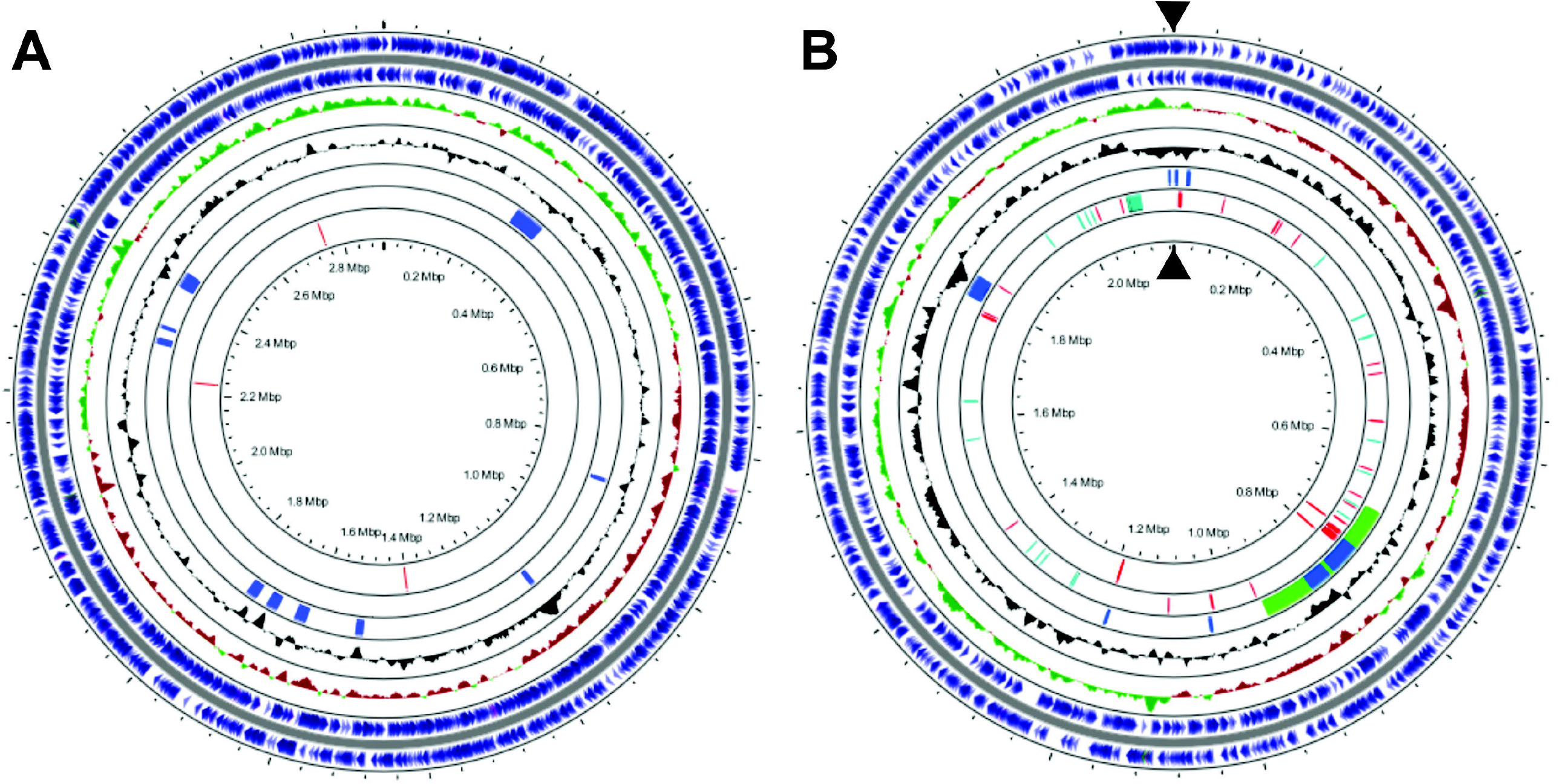
Circular representation of *A. tumefaciens* 6N2 genome. In the circular (A) chromosome, from outer to inner are represented CDS in each strand (blue), GC skew (green and red), GC content (black), genomic islands (blue) and the luxR orthologs *rhiR, atxR* and *solR* (red, clockwise sense). In the linear (B) chromosome, are represented CDS in each strand (blue), GC skew (green and red), GC content (black), genomic islands predicted with Islanviewer and ICEs predicted with ICEfinder (blue and green), T4SS and T6SS (red and cyan), and the tQS, QS2 and QS1 (red, clockwise sense). Black triangles indicate the extreme of the linear chromosome.

### Identification of quorum sensing systems in *A. tumefaciens* 6N2

Strain 6N2 genomic sequence showed the absence of a QS system comparable to the TraI/TraR QS system of *A. fabrum* strain C58 [3]. A more complex architecture was identified in the linear chromosome (Fig. 3B and Fig. 4). A first system, here named QS1 (coordinates 1,189,496-1,191,920) was composed of *luxR* orthologues AT6N2_L1344 and AT6N2_L1347, one overlapped by the last 4 bp of the *luxI* ortholog AT6N2_L1345. Considering this R-IR topology, similar to *A. fabacearum* strain P4 QS system [13], genes were named accordingly *cinR, cinI* and *cinX*.

**Figure 4.**
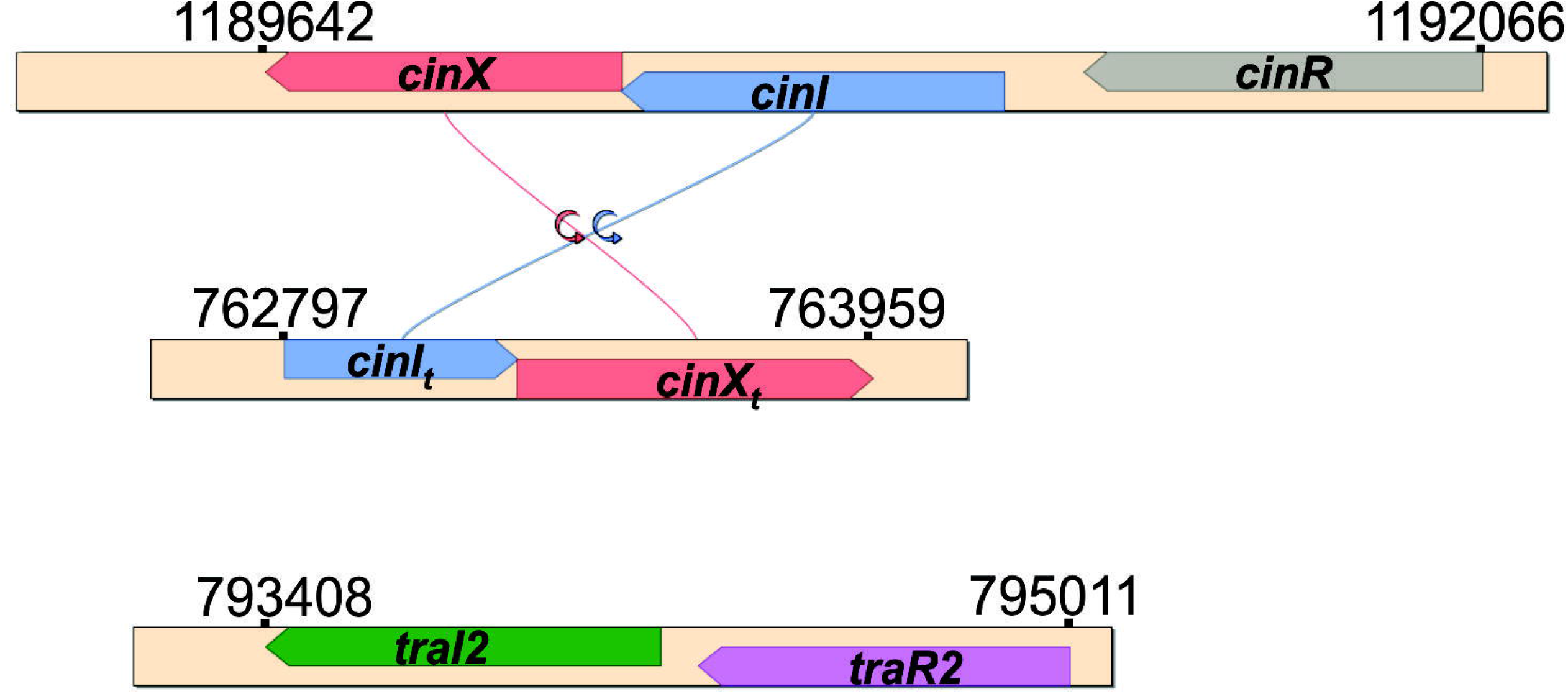
Architecture and topology of *A. tumefaciens* 6N2 QS systems based on AHL signals. In the linear chromosome, QS1 composed of *cinR, cinI* and *cinX*, and QS2 composed of *traI2* and *traR2* were identified. A truncated tQS system composed of *cinX* and a truncated version of *cinI* apparently arose from a partial duplication and inversion of QS1. Respective coordinates are shown. Figure was prepared with SimpleSinteny software.

A second QS system, named QS2 (coordinates 793,262-794,901), was found in the linear chromosome transcribed in the same direction as QS1 (Fig. 3B and Fig. 4). With a R-I topology, QS2 was composed of the *luxI* and a *luxR* orthologues *traI2* (AT6N2_L0888) and *traR2* (AT6N2_L0889), respectively. A truncated system, here named tQS (coordinates 762,651-763,813) was also found in the linear chromosome and in the opposite direction to QS1 and QS2 (Fig. 3B and Fig. 4). tQS, composed of a *luxR* (AT6N2_L0841) and a truncated *luxI* (AT6N2_L0840) orthologues, was probably originated from a partial duplication and inversion of QS1. Indeed, *luxR* and *cinX* showed 90% identity (641/711); *luxI* and *cinI* showed 92% identity (420/456). With 456 nucleotides, this *luxI* is significantly shorter than *cinI* (765 nucleotides). tQS genes were named accordingly as *cinX*_t_ and *cinI*_t_ (Fig. 3B and Fig. 4). Three *luxR* orthologues were identified in the circular chromosome (Fig. 3A). AT6N2_C1772 (coordinates 1,401,123-1,400,383) was named *rhiR* for its homology with *A. radiobacter rhiR* and *A. fabrum* C58 Atu0707. AT6N2_C2737 (coordinates 2,231,916-2,232,653) was named *solR* for its homology with *A. radiobacter solR* and *A. fabrum* C58 Atu2727. AT6N2_C3352 (coordinates 2,749,807-2,750,523) was named *atxR* due to its homology with *A. fabrum* C58 *atxR* (Atu2285) (Fig. 3A). The analysis of the putative aminoacid sequences of AtxR, SolR and RhiR showed the characteristics domains for DNA and autoinducer binding (data not shown).

A search in *Agrobacterium* genomes allowed the identification of strains with similar topologies in the QS systems. *A. tumefaciens* strain 5A, *A. fabacearum* P4, *A. deltaense* strains RV3 and NCPPB1641 and *A. radiobacter* strain DSM30147 exhibit QS systems similar to QS1 (R-IR topology). Synteny throughout 16,200 bp upstream QS1 is highly conserved among these strains (Suppl. Fig. 2A). QS2 topology (R-I) was detected in *A. tumefaciens* strains S2, S33, *Agrobacterium* sp. strain SUL3 and *A. arsenijevicii* strain KFB330, but with no conservation of synteny (data not shown). Homologues of *atxR* (Suppl. Fig. 2B), *solR* (Suppl. Fig. 2C) and *rhiR* (Suppl. Fig. 2D) were identified in all these strains, including strain C58, with synteny highly conserved, encompassing 235,500 bp, 785,000 bp, and 98,000 bp, respectively.

A multiple alignment of aminoacid sequences of LuxI orthologues showed identities higher between strain 6N2 CinI and proteins with the same R-IR topology (Suppl. Fig. 3A). Orthologues with R-I topology like 6N2 TraI2 presented less similarity among them. CinR, CinX, CinXt, AtxR, SolR and RhiR also showed high similarities with orthologues sharing the topology and synteny (Suppl. Fig. 3B). Similar to TraI2, low similarities were found among 6N2 TraR2 and orthologues with the R-I topology. To note, all the 6N2 LuxI and LuxR orthologues showed low similarities with *A. fabrum* C58 TraI and TraR.

### Modulation of *A. tumefaciens* strain 6N2 proteome by QS

Quorum quenching strategy with pME6863 [14] was successful for the attenuation of the *A. tumefaciens* 6N2 (see Suppl. Figure 4). At late exponential growth phase, no growth differences were found between *A. tumefaciens* 6N2 carrying the empty control vector pME6000 and *A. tumefaciens* 6N2 (pME6863). Both strains attained cell densities of ~ 1.5 10^9^ CFU ml^-1^ (data not shown). A total of 2,637 proteins were identified in extracts from single cultures of *A. tumefaciens* 6N2 (pME6000) and *A. tumefaciens* 6N2 (pME6863). Considering a *p*≤0.05 and a FC ≥ 1.5, the attenuation of the QS activity altered the relative abundances of 64 proteins in single cultures of *A. tumefaciens* strain 6N2 (6N2^QSPR^ group), coded in the circular (37) and the linear (27) chromosome (Fig. 5A and Table 1). Thirty-three were more abundant in *A. tumefaciens* strains 6N2 (pME6000) in comparison with *A. tumefaciens* strain 6N2 (pME6863), indicating an upregulation by QS (6N2^QSPR^_up_ subgroup); 31 in 6N2^QSPR^ group were less abundant in *A. tumefaciens* 6N2 (pME6000), indicating a downregulation by QS (6N2^QSPR^_dw_ subgroup) (Fig. 5A and Table 1).

**Figure 5.**
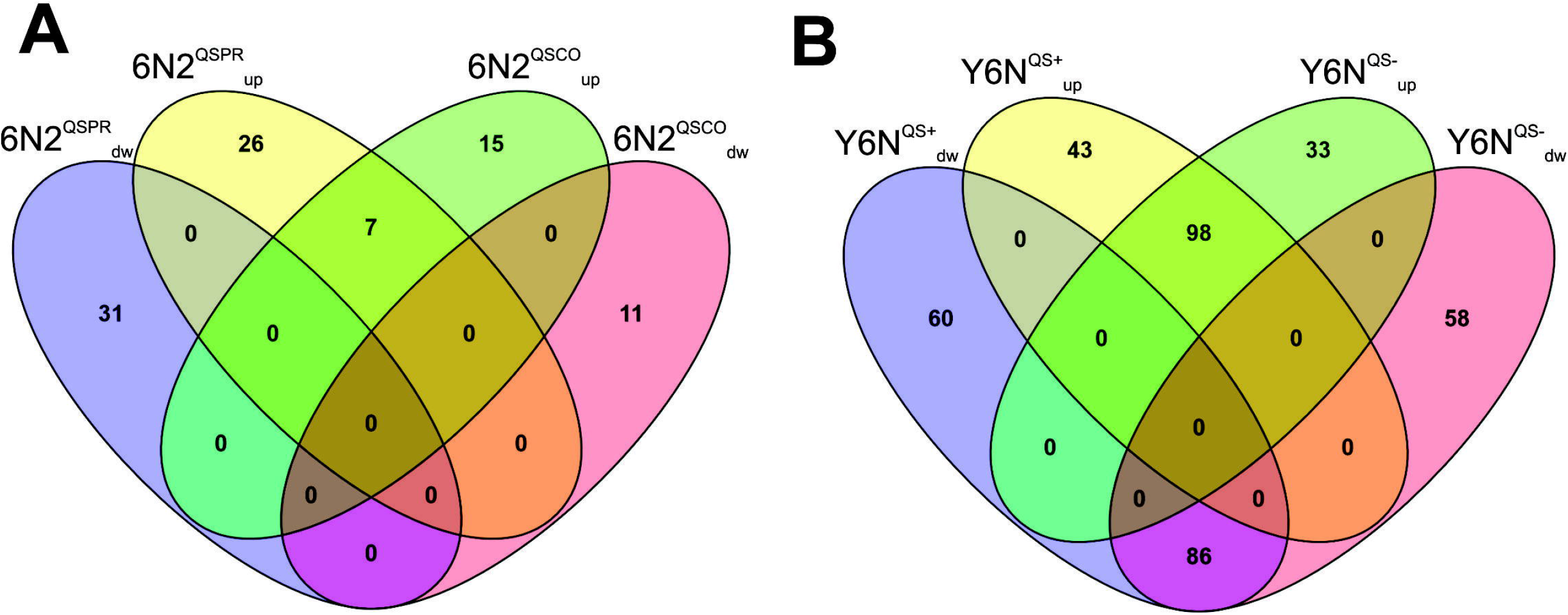
Summary of proteomic analysis of *A. tumefaciens* 6N2 and *M. guilliermondii* 6N. Venn diagrams shows bacterial (A) and yeast subgroups of proteins. In 6N2, 7 common AHL-based QS-regulated proteins were found in the 6N2^QSPR^_up_ and 6N2^QSCO^_up_ subgroups. In the yeast, 184 differentially accumulated proteins (98+86) were attributed to the presence of the bacterium, with independence of the QS activity. Venn diagram was prepared with Venny web program (https://bioinfogp.cnb.csic.es/tools/venny/index.html).

6N2^QSPR^ proteins were classified in eggNOG, mainly in Energy production and conversion (4), and Amino acid transport and metabolism (8); 14 were classified as Function unknown (Suppl. Fig. 5A and 5B). To gain insight into the influence of QS on *A. tumefaciens* strain 6N2 physiology, the ontology of 6N2^QSPR^ group proteins were analyzed (Suppl. Fig. 6). In the Biological Process (BP) ontology of 6N2^QSPR^ group (Suppl. Fig. 6A and 6B), most were classified in Biosynthesis (GO:0009058), Cell organization and biogenesis (GO:0016043), Metabolism (GO:0008152), Transport (GO:0006810), and Nucleobase, nucleoside, nucleotide and nucleic acid metabolism (GO:0006139). The Cellular Component (CC) ontology (Suppl. Fig. 6C and 6D) showed most of the proteins classified in Cell (GO:0005623), and Intracellular (GO:0005622). In the Molecular Function (MF) ontology (Suppl. Fig. 6E and 6F), most of the 6N2^QSPR^ group proteins were in Binding (GO:0005488), Catalytic activity (GO:0003824), Hydrolase activity (GO:0016787) and Transferase activity (GO:0016740).

Regulatory and signaling proteins were identified in the 6N2^QSPR^ group: CinX regulatory protein (AT6N2_L1344), LacI-type regulator (AT6N2_C1926), and GntR-type (AT6N2_C0879) transcriptional regulators in 6N2^QSPR^_up_ subgroup; two sensor histidine kinases (AT6N2_C3453 and AT6N2_C3125), and YebC-like regulator (AT6N2_L1564) in 6N2^QSPR^_dw_. Several proteins in 6N2^QSPR^ group were related to transport of small molecules or ions: a mechanosensitive ion channel protein (AT6N2_C0650), a DMT family transporter (AT6N2_C0483), an ABC transporter permease (AT6N2_C3519), a multidrug efflux RND transporter permease subunit (AT6N2_C3101) and an ABC transporter substrate-binding protein (AT6N2_L1359) in 6N2^QSPR^_up_ subgroup; a transporter substrate-binding domain-containing protein (AT6N2_C3262), a dicarboxylate/amino acid:cation symporter (AT6N2_L0298), an ABC transporter substrate-binding protein (AT6N2_L0602) and an ABC transporter ATP-binding protein/permease (AT6N2_L1331) in 6N2^QSPR^_dw_.

Several proteins in 6N2^QSPR^_up_ can be highlighted. Orthologues of AT6N2_L0856 (pilus assembly protein), AT6N2_L0857 (Conjugal Transfer Protein D), and AT6N2_L2010 (Mobilization Protein C) are related to the QS-regulated transfer of pTi and pAt in strain C58, and of pAt in strain P4 through a type IV secretion system. In addition to CinX, RhiR (AT6N2_C1772) was the only protein from the complex 6N2 QS system identified in the proteomic analysis. RhiR was over accumulated (p<0.05) when the QS system was attenuated with a FC=1.49, just below the arbitrary limit established in this work.

### The yeast *M. guilliermondii* 6N alters the QS regulation in *A. tumefaciens* 6N2

At late exponential growth phase, cell densities of both *A. tumefaciens* 6N2 (pME6000) and *A. tumefaciens* 6N2 (pME6863) were one log unit lower than in pure cultures (~ 3.6 10^8^ CFU ml^-1^), with no differences between the two strains (data not shown). A total ion current normalization based exclusively on bacterial proteins were applied to allow a comparison between pure and dual cultures.

A notable reduction in QS-regulated proteins was determined in dual culture with *M. guilliermondii* 6N. Only 33 proteins (6N2^QSCO^ group) were influenced by the QS activity, which were coded in the circular (19) and the linear (14) chromosome (Table 1). In 6N2^QSCO^, 22 were more abundant in strain 6N2 (pME6000), indicating an upregulation by QS (6N2^QSCO^_up_ subgroup) in co-culture. Eleven proteins of 6N2^QSCO^ were less abundant, indicating a downregulation (6N2^QSCO^_dw_ subgroup) in dual culture (Fig. 5A and Table 1). 6N2^QSCO^ proteins were mainly classified (Suppl. Fig. 5A and 5B) in Transcription (3) and Function unknown (12). In BP ontology (Suppl. Fig. 6A and 6B), most were classified in Biosynthesis (GO:0009058), and Metabolism (GO:0008152). In CC ontology (Suppl. Fig. 6C and 6D), the majority were classified in Cell (GO:0005623) and Plasma membrane (GO:0005886). The MF ontology of 6N2^QSCO^ (Suppl. Fig. 6E and 6F) showed most classified in Binding (GO:0005488), Catalytic activity (GO:0003824), and Hydrolase activity (GO:0016787).

In the 6N2^QSCO^ group, 4 were regulatory proteins or related to signal transduction: CinX (AT6N2_L1344), ArsR family transcriptional factor (AT6N2_C3363) and Xre family transcriptional factor (AT6N2_L0663) in the 6N2^QSCO^_up_ subgroup; a Response regulator PleD (AT6N2_C1017) in 6N2^QSCO^_dw_. Four were related to transport of nutrients: a component of a metal ABC transporter permease (AT6N2_C1510) and a component of a zinc ABC transporter (AT6N2_C0769) in 6N2^QSCO^_up_ subgroup; a sugar ABC transporter ATP-binding protein (AT6N2_L0645) and an ABC transporter substrate-binding protein (AT6N2_L1359) in 6N2^QSCO^_dw_. Phage proteins were identified in the 6N2^QSCO^_dw_ subgroup: a major capsid protein (AT6N2_C0409), an ATP-binding protein (AT6N2_C0382) and a DNA polymerase III subunit beta (AT6N2_C0386), all part of the predicted prophage *16-3*. The comparison between the different subgroups showed only 7 common proteins between 6N2^QSPR^_up_ and 6N2^QSCO^_up_ (Fig. 8A): Nucleotidyltransferase (AT6N2_L0014), Hypothetical Protein (AT6N2_L0851), Pilus assembly protein (AT6N2_L0856), Conjugal Transfer Protein D (AT6N2_L0857), CinX (AT6N2_L1344), TauD/TfdA family dioxygenase (AT6N2_L1355) and ABC transporter substrate-binding protein (AT6N2_L1359). No common proteins were found in the comparison between 6N2^QSPR^_dw_ and 6N2^QSCO^_dw_ (Fig. 8A). Similar to single cultures, CinX and RhiR were the only components of the 6N2 QS system identified, though in co-culture RhiR was not supported statistically (p>0.05).

### The QS activity of *A. tumefaciens* 6N2 modifies the proteome of *M. guilliermondii* 6N

The yeast *M. guilliermondii* 6N reached a cell density of ~ 1.2 10^8^ CFU ml^-1^ in pure culture, one log unit higher in comparison to dual cultures with *A. tumefaciens* 6N2 (pME6000) (3.2 10^7^ CFU ml^-1^) and *A. tumefaciens* 6N2 (pME6863) (4.6 10^7^ CFU ml^-1^). Similar to *A. tumefaciens* 6N2, a total ion current normalization based exclusively on yeast proteins were applied to allow a comparison between pure and dual cultures.

The comparison of the *M. guilliermondii* proteomes between pure and dual cultures, showed 287 proteins (Table 2) whose abundances were modified by *A. tumefaciens* (pME6000) (Y6N^QS+^ group): 141 upregulated (Y6N^QS+^_up_ subgroup) and 146 downregulated (Y6N^QS+^_dw_ subgroup). On the other hand, 275 proteins (Table 2) were modified by *A. tumefaciens* (pME6863) (Y6N^QS-^ group): 131 up-accumulated (Y6N^QS-^_up_ subgroup) and 144 down-accumulated (Y6N^QS-^_dw_ subgroup) (Figure 5B). To note, 98 proteins were common among Y6N^QS+^_up_ and Y6N^QS-^_up_ subgroups; 86 were common among Y6N^QS+^_dw_ and Y6N^QS-^_dw_ (Figure 5B and Table 2). These 184 (98+86) common proteins were then attributed to the presence of the bacterium, independently of the agrobacterial QS activity, and in consequence no longer considered in this report. In comparison to the pure culture, among the fungal proteins increased due to strain 6N2 QS activity, 43 were identified in Y6N^QS+^_up_ subgroup and 33 were in Y6N^QS-^_up_. The categories of each subgroup in eggNOG were dissimilar (Suppl. Fig. 7A). For instance, several categories were more numerous in Y6N^QS+^_up_, including RNA Processing and modification, Energy production and conversion, Amino acid transport and metabolism, Lipid transport and metabolism, Posttranslational modification, protein turnover, chaperones, Secondary metabolites biosynthesis, transport and catabolism, and Intracellular trafficking, secretion, and vesicular transport. In Y6N^QS+^_dw_, Translation, ribosomal structure and biogenesis was more numerous (Figure 7A). Biological Process (BP), Cellular Component (CC) and Molecular Function (MF) ontologies showed differences among Y6N^QS+^_up_ and Y6N^QS-^_up_. These dissimilarities were more notorious in Biosynthesis (GO:0009058), Catabolism (GO:0009056), Metabolism (GO:0008152) and Protein metabolism (GO:0019538) of BP ontology; Cell (GO:0005623) and Intracellular (GO:0005622) of CC ontology; and Binding (GO:0005488), Catalytic activity (GO:0003824), Nucleic acid binding (GO:0003676), Nucleotide binding (GO:0000166) and Transporter activity (GO:0005215) of MF ontology (Suppl. Figures 8A, 8C and 8D). In Y6N^QS+^_up_ subgroup, it is to highlight the identification of E3 ubiquitin-protein ligase (A5DGJ2), E2 ubiquitin-conjugating enzyme (A5DL67), protein kinase domain-containing protein (A5DE57) and RAS-domain containing protein (A5DKQ9). In Y6N^QS-^_up_, Vacuolar proton pump subunit B (A5DEC0), Vacuolar protein sorting-associated protein (A5DHU0), V-type proton ATPase subunit (A5DLL8) and Phosphoenolpyruvate carboxykinase (A5DD88). The same observation was made in the comparison of proteins down-regulated by the agrobacterial QS activity in Y6N^QS+^_dw_ and Y6N^QS-^_dw_. Data from eggNOG (Suppl. Fig. 7B) showed, for instance, Y6N^QS+^_dw_ proteins more numerous in categories that include RNA Processing and modification, Coenzyme transport and metabolism, and Transcription. Proteins in Posttranslational modification, protein turnover, chaperones were more numerous in Y6N^QS-^_dw_ than in Y6N^QS+^_dw_.

Biological Process (BP), Cellular Component (CC) and Molecular Function (MF) ontologies of Y6N^QS+^_dw_ and Y6N^QS-^_dw_ also presented differences in the values of proteins assigned to each category (Suppl. Fig. 8B, 8D and 8F). Main differences were in Cell communication, Cell cycle, Cell organization and biogenesis, Organelle organization and biogenesis, Protein metabolism, Cell, Intracellular, Catalytic activity and Transferase activity, among others.

## Discussion

Strain 6N2 belongs to the group of avirulent and commensal agrobacteria. This strain was obtained from sugarcane, which is in contrast to dicots is not susceptible to crown gall formation [15]. An At plasmid is also absent in its genome, indicating that strain 6N2 is a plasmid-less agrobacterium. Possibly, this particular niche, with no selective pressure to maintain extrachromosomal replicons, had molded the 6N2 genome [16].

In comparison with strain C58 [3], strain 6N2 produces four AHLs, and two AHL synthases are encoded in its linear chromosome. One of this molecule, 3OHC8-HSL, has also been reported in the non-pathogenic strain P4 [13], which similarly harbors CinI coded in a QS system with the same R-IR topology as 6N2 QS1. It is then plausible that 6N2 CinI is also involved in 3OHC8-HSL production. It is tempting to assign the synthesis of 3OHC10, 3OHC12-HSL and 3OC12-HSL to 6N2 TraI2. It has to be considered that an enzyme can be involved in the production of more than one AHL [17]. To date, the only *Agrobacterium* LuxI homolog characterized with a QS2 architecture is *A. vitis* AvsI, involved in the production of multiple long chain-AHLs [18]. AtxR, SolR and RhiR are also present in strain C58 [19], though their role in QS have not been evaluated. AviR, a SolR homolog, is a key regulator of the pathogenesis in *A. vitis* [20,21]. A number of three LuxR orphans (i.e., LuxR homologs unpaired to LuxI homologs) in *A. tumefaciens* 6N2 is comparable to some of the strains mentioned in this manuscript (2 LuxR orphans in *A. arsenijevicii* KFB330; 3 in *A. tumefaciens* S2, *A. tumefaciens* S33, *A. radiobacter* DSM30147 and *A. fabrum* C58; 4 in SUL3, *A. fabacearum* P4 and *A. deltaense* RV3; 5 in *A. tumefaciens* 5A; 6 in *A. deltaense* NCPPB1641).

The lack of similar mechanisms to 6N2 QS1 in linear chromosomes of other agrobacteria, could be associated to the plasticity of these replicons. Also tQS is in a regions of genome plasticity and predicted in an ICE element. This fact could be related to the truncated nature of *cinI*_t_. It is plausible that this truncated QS system is a remnant of a duplication and inversion event, without activity. Additionally, the putative CinI_t_ is only 151 residues long, meaning a lack of 103 residues in the *N*-terminus in comparison to CinI. However, a truncated *luxI* homolog in *Methylobacterium extorquens* AM1 [22], controls the AM1 QS systems.

Considering that agrobacterial QS systems are usually involved in plasmid conjugation [13,23,24], their localization in the linear chromosome of strain 6N2 open the question about their functions. Several 6N2 proteins regulated by QS are related to conjugative functions: Pilus assembly protein, Conjugal Transfer Protein D and Mobilization Protein C. It is plausible that the 6N2 QS activity is involved in the conjugation of other genetic elements, though they could also be remnants of an integration event of a conjugative plasmid. The C58 linear chromosome also harbors homologs suggested to participate in the mobilization of part of the chromosome [25]. To note, a mobile element of 228,159 bp (coordinates 712,734-940,892), encompassing tQS, QS2 and the T4SS is predicted in the linear chromosome. It is possible that 6N2 QS systems also modify the bacterial metabolism, considering the proteins under the influence of the QS activity, related to energy production and conversion, amino acid transport and metabolism, transport of ions and small molecules. It remains to be elucidated whether this regulation is exerted directly through the LuxR homolog(s), or through other regulatory proteins found regulated by the 6N2 QS activity.

The finding that *M. guilliermondii* 6N alters the bacterial proteome is not surprising. It is now clear that the co-cultivation of different species activates gene clusters otherwise silenced, and vice versa, a process driven by chemical and physical interactions [26]. Most astonishing is the modification of the 6N2 QS regulation in co-culture with the yeast: several proteins remain regulated by QS independently of *M. guilliermondii* 6N, but others are affected by the yeast. The 7 common proteins in 6N2^QSCO^_up_ and 6N2^QSPR^_up_ could be attributed to a direct QS regulation, while the others could be indirect or susceptible of modification by an environmental factor like the presence of the yeast. To note, three of these common proteins (Hypothetical protein AT6N2_L0851, Pilus assembly protein AT6N2_L0856 and Conjugal Transfer Protein D AT6N2_L0857) are coded between QS2 and tQS. As mentioned before (see Modulation of *A. tumefaciens* strain 6N2 proteome by QS), the Pilus assembly protein and the Conjugal Transfer Protein D have been related to the conjugal transfer of pAt in *A. tumefaciens* strain P4. Other three of them (Autoinducer binding domain-containing protein CinX AT6N2_L1344, TauD/TfdA family dioxygenase AT6N2_L1355 and ABC transporter substrate-binding protein AT6N2_L1359) are coded close to QS1.

This accompanying microorganism could degrade, metabolize or modify the QS signals modulating in consequence the QS activity [27]. However, it is unlikely that the modification in the 6N2 QS regulation can be attributed to a fungal inactivation of QS signals. Even though QQ is prevalent in yeasts, *M. guilliermondii* 6N exhibits only a weak capacity for inactivating AHLs [10]. Probably other mechanisms take part in the *M. guilliermondii* 6N-*A. tumefaciens* 6N2 interactions and the subsequent alteration of the QS regulation, as described in oral biofilms, where cell-cell contacts and production or depletion of metabolites intervene in the establishment of microbial communities [28]. Indeed, some QS-regulated proteins are related to the transport and metabolism of ions and metabolites, as mentioned above. A recent report presented a model showing how an environmental cue, through dedicated regulators, act on QS signals or signal receptors modulating the gene expression [29]. Particular attention deserve the prophage *16-3* proteins identified in 6N2^QSCO^_dw_ subgroup, since this is in concordance with the “piggyback-the-winner” theory, which predicts a lysogenic switching at high cell densities [30]. The relationship between QS and lysogeny has been proven for the induction of the lytic cycle of ΦH2O [31]. Our proteomic results suggest that, in addition to QS, other environmental factors, like the simultaneous presence of other microorganism, could influence the phage cycle.

First described in the *Pseudomonas aeruginosa*-*Candida albicans* interactions, it is now clear that QS molecules not only influence the physiology of the signaling microorganism but also that of surroundings microorganisms [32,33]. In this work, we describe for the first time the alteration of a yeast proteome by the bacterial QS activity. It is probable that the AHLs, absent or strongly diminished in the co-culture with *A. tumefaciens* (pME6863), have a direct effect on the yeast. Although no AHL receptor has been described in eukaryotic cells, these molecules can interact with biological membranes modifying the dipole potential [34]. An indirect mechanism is also possible for this modification of the fungal proteome: a QS-mediated alteration of the bacterial physiology could modify the profile of metabolites in the culture medium, altering the fungal proteome. Both direct and indirect mechanisms are not mutually exclusive. Beyond the mechanism that modulates the yeast proteome, it is to note that relevant events are being modified. For instance, a RAS domain-containing protein (A5DKQ9), an E3 ubiquitin-protein ligase (A5DL67) and an E2 ubiquitin-conjugating enzyme (A5DL67) in Y6N^QS+^_up_ subgroup, and an USP domain-containing protein (A5DNC6) and protein FYV10 (A5DFE2) in Y6N^QS+^_dw_ suggest an up- regulation of an ubiquitylation process, less prevalent when the QS activity is attenuated [35,36]. In agreement, protein metabolism is one of the main terms in BP ontology showing differences between Y6N^QS+^_up_ and Y6N^QS-^_up_. In contrast, a vacuolar proton pump subunit B (A5DEC0), a vacuolar protein sorting-associated protein (A5DHU0) and a V-type proton ATPase subunit (A5DLL8) identified in Y6N^QS-^_up_, together with the the GO term “vacuole” in CC ontology put the focus in this organelle, key compartment in the fungal cell [37]. Though focused in an *in vitro* description, our results indicate the importance of the *in planta* characterization of the *A. tumefaciens* 6N2 QS system, for evaluating its ecological and physiological relevance, including its role in growth and survival. The complete elucidation of the mechanism beneath the *A. tumefaciens* 6N2-*M. guilliermondii* 6N interactions requires the consideration of the QS-influenced proteins, those guided by the presence of the second microorganism and, importantly, also those whose abundances are constant. However, results presented in this report allow a first insight to the complexity of the interactions between these two microorganisms.

## Materials and methods

### Microorganisms and growth conditions

*A. tumefaciens* 6N2 and *M. guilliermondii* 6N were cultured at 30 ºC in nutrient broth (NB) (peptone 5 g L^-1^; yeast extract 3 g L^-1^). *Escherichia coli* DH5α harboring plasmids pME6000 [38] or pME6863 [14] were cultured in Luria Bertani broth at 37 ºC. When required, media were supplemented with agar, 15 g L^-1^, ampicillin 100 µg mL^-1^, tetracycline 15 µg mL^-1^ or cycloheximide 50 µg mL^-1^.

### AHL identification

Five hundred mL of NB broth were inoculated with an overnight culture of *A. tumefaciens* 6N2, and incubated aerobically at 30 ºC for 24 h until late exponential growth phase. Supernatants were extracted twice with acidified ethyl acetate [39]. Concentrated extracts were analyzed by UPLC/ESI MS/MS (Waters Aquity UPLC-TQD) with an Acquity HSS C18 (2.1 mm × 50 mm; 1.8 μm) at 20 ºC with a flow of 0.6 mL min^-1^ and a gradient of 10% acetonitrile with 0.1% formic acid to 100% acetonitrile with 0.1% formic acid in 5 min as mobile phase. AHL identifications were performed by comparison of fragmentation patterns with those of commercial AHLs [39].

### Genomic sequencing and annotation

*A. tumefaciens* 6N2 genomic DNA was extracted from a 10 mL overnight culture. Genome sequence was obtained utilizing single-molecule real-time sequencing technology (Pacific Biosciences) (see Supp. Materials for details). Annotation was performed with the MicroScope platform [40] and BASys [41]. For the identification of QS genes, BLAST searches were performed on strain 6N2 genome utilizing as query the *traI* and *traR* of *A. fabrum* C58 and related microorganisms (see Supp. Materials for details). Sequences were deposited in Genbank under accession numbers CP072308 and CP072309.

### Attenuation of QS activity

A quorum quenching (QQ) strategy was developed, using the vector pME6863 [14] that allows the constitutive expression of the *Bacillus* spp. AiiA lactonase. The vector was conjugated from DH5α (pME6863) into strain 6N2 in a triparental mating with *E. coli* DH5α (pRK2013) [42] on LB agar plates for 24 h at 30 ºC. pME6000 [38] was independently conjugated as negative control. Transconjugants were selected on LB agar supplemented with ampicillin and tetracycline. To confirm the QQ strategy, organic extracts were analyzed by RP-TLC using *A. tumefaciens* NT1 (pZLR4) as bioreporter strain [43,44].

### Pathogenicity assays

Crown gall tumor formation was assessed on tomato and *Arabidopsis thaliana* plants. *A. tumefaciens* strain 6N2 was cultured on NB agar for 48 h, cells were aseptically scraped off and resuspended in sterile water at a final density of 10^7^ CFU mL^-1^. Cell suspension was inoculated in 4-cm cuts between the first and second node on the stems of young tomato plants. *A. thaliana* was inoculated below the first node. Plants were incubated 2 weeks at 25 °C under 16 h illumination and inspected for the apparition of tumors. *A. fabrum* C58 and sterile water were utilized as positive and negative controls.

### Preparation of protein extracts and proteomic analysis

Two hundred and fifty mL flasks containing 20 mL of NB broth were inoculated at an initial concentration of ~ 10^7^ CFU mL^-1^ for *A. tumefaciens* 6N2 (pME6000) or *A. tumefaciens* 6N2 (pME6863), and ~ 10^6^ CFU mL^-1^ for *M. guilliermondii* 6N. Dual cultures of *A. tumefaciens* 6N2 (pME6000) plus the yeast, and *A. tumefaciens* 6N2 (pME6863) plus the yeast, were prepared with those cell densities. Flasks were incubated aerobically at 30 ºC for 24 h until late exponential growth phase. Protein extracts were obtained using the YPX extraction kit (EXPEDEON), and concentrations were determined with the QuantiPro BCA (SigmaAldrich). Three independent samples were analyzed for each pure or mixed culture. Protein samples were trypsin digested and peptide mixtures were analyzed by a Q-Exactive mass spectrometer coupled to an Easy-nLC system (both from Thermo Scientific). All MS/MS data were processed with Proteome Discoverer 2.1 (Thermo Scientific) coupled to an in-house Mascot search server (Matrix Science, Boston, MA; version 2.5.1). Proteins showing a fold change (FC) ≥ 1.5 and an ANOVA p≤0.05 were considered as differentially accumulated (see Supp. Materials for details). Complete datasets are available in the ProteomeXchange Consortium via the PRIDE [45] partner repository with the identifier PXD025730.

## Acknowledgement

This work was supported by the Consejo Nacional de Investigaciones Científicas y Técnicas (CONICET, PIP 2015 N°0946, PU-E22920160100012CO), Agencia Nacional de Promoción Científica y Tecnológica (PICT 2016 Nº 0532; PICT 2016 N° 2013), and Secretaría de Ciencia, Arte e Innovación Tecnológica from the Universidad Nacional de Tucumán (PIUNT D609). Kok-Gan Chan thanks the financial support from the University of Malaya (FRGS grant no. FP022-2018A). Carlos Nieto-Peñalver thanks the support from Université Paris Diderot through the Alicia Moreau Chair.

---

Suppl. Figure 1. Acyl homoserine lactones produced by *A. tumefaciens* 6N2. UPLC/ESI MS/MS analysis allowed the identification of *N*-3-hydroxy-octanoyl-homoserine lactone (OHC8-HSL), *N*-3-hydroxy-decanoyl-homoserine lactone (OHC10-HSL), *N*-3-oxo-dodecanoyl-homoserine lactone (OC12-HSL) and *N*-3-hydroxy-dodecanoyl-homoserine lactone (OHC12-HSL).

Suppl. Figure 2. Conservation of synteny around the *A. tumefaciens* 6N2 QS systems related to AHLs. Synteny upstream QS1 (A) spans 16200 bp and is shared by the *Agrobacterium* strains 5A, P4, RV3, NCCPB1641 and DSM30147. Syntenies around *atxR* (B), *solR* (C) and *rhiR* (D) span 235500 bp, 785000 bp, and 98000 bp, respectively, and can be identified in all the strains analyzed. Due to the length of the sequences, only part of the synteny is shown. Respective 6N2 coordinates are shown. Analysis were performed with SimpleSinteny software.

Suppl. Figure 3A. Similarities among *Agrobacterium* LuxI orthologs. 6N2 orthologs are indicated with the corresponding protein name. Orthologs of other strains are indicated with the corresponding Genbank accession number. Identity matrix constructed with BioEdit was visualized as a heatmap with MORPHEUS. Identity values are indicated in each square. Low identities are represented with orange-red shades; high identities are with green shades; medium identities are in yellow (see scale bar).

Suppl. Figure 3B. Similarities among *Agrobacterium* LuxR orthologs. 6N2 orthologs are indicated with the corresponding protein name. Orthologs of other strains are indicated with the corresponding Genbank accession number. Identity matrix constructed with BioEdit was visualized as a heatmap with MORPHEUS. Identity values are indicated in each square. Low identities are represented with orange-red shades; high identities are with green shades; medium identities are in yellow (see scale bar).

Suppl. Figure 4. Attenuation of QS activity in *A. tumefaciens* 6N2. Concentrated extracts of *A. tumefaciens* 6N2 (pME6000) and *A. tumefaciens* 6N2 (pME6863) were analyzed by RP-TLC utilizing MeOH:H2O (6:4) as mobile phase. 3OHC8-HSL (4 pmol), 3OHC10-HSL (0.25 nmol) and 3OHC12-HSL (2.5 nmol) were utilized as standards (Stds).

Suppl. Figure 5. Functional classification of *A. tumefaciens* 6N2 QS regulated proteins. 6N2^QSPR^_up_ and 6N2^QSCO^_up_ subgroups are compared in (A). 6N2^QSPR^_dw_ and 6N2^QSCO^_dw_ subgroups are compared in (B). In each figure, full bars correspond to pure cultures and dashed bars correspond to cocultures. eggNOG database was utilized for the analysis. C, Energy production and conversion; E, Amino acid transport and metabolism; F, Nucleotide transport and metabolism; G, Carbohydrate transport and metabolism; H, Coenzyme transport and metabolism; I, Lipid transport and metabolism; J, Translation, ribosomal structure and biogenesis; K, Transcription; L, Replication, recombination and repair; O, Posttranslational modification, protein turnover, chaperones; P, Inorganic ion transport and metabolism; Q, Secondary metabolites biosynthesis, transport and catabolism; S, Function unknown; T, Signal transduction mechanism; U, Intracellular trafficking, secretion, and vesicular transport; NA, not assigned.

Suppl. Figure 6. Ontology analysis of differentially accumulated proteins of *A. tumefaciens* 6N2. Figures show the number of bacterial proteins regulated by the QS activity, associated to the respective GO terms. BP (upper), CC (middle) and MF (lower) ontologies of bacterial proteins are shown. In each figure, full bars correspond to pure cultures and dashed bars correspond to cocultures. 6N2^QSPR^_up_ and 6N2^QSCO^_up_ are depicted in green (A). 6N2^QSPR^_dw_ and 6N2^QSCO^_dw_ are depicted in red (B).

Suppl. Figure 7. Functional classification of *M. guilliermondii* 6N proteins regulated by 6N2 QS activity. Y6N^QS+^_up_ and Y6N^QS-^_up_ subgroups are compared in (A). Y6N^QS+^_dw_ and Y6N^QS-^_dw_ subgroups are compared in (B). In each figure, full bars correspond to cocultures with *A. tumefaciens* 6N2 (pME6000) and dashed bars correspond to cocultures with *A. tumefaciens* 6N2 (pME6863). eggNOG database was utilized for the analysis. A, RNA Processing and modification; B, Chromatin structure and dynamics; C, Energy production and conversion; D, Cell cycle control, cell division, chromosome partitioning; E, Amino acid transport and metabolism; F, Nucleotide transport and metabolism; G, Carbohydrate transport and metabolism; H, Coenzyme transport and metabolism; I, Lipid transport and metabolism; J, Translation, ribosomal structure and biogenesis; K, Transcription; L, Replication, recombination and repair; M, Cell wall/membrane/envelope biogenesis; O, Posttranslational modification, protein turnover, chaperones; P, Inorganic ion transport and metabolism; Q, Secondary metabolites biosynthesis, transport and catabolism; S, Function unknown; T, Signal transduction mechanism; U, Intracellular trafficking, secretion, and vesicular transport; Y, Nuclear structure; Z, Cytoskeleton; NA, not assigned.

Suppl. Figure 8. Ontology analysis of differentially accumulated proteins of *M. guilliermondii* 6N. Figures show the number of yeast proteins influenced by the *A. tumefaciens* 6N2 QS activity, associated to the respective GO terms. BP (upper), CC (middle) and MF (lower) ontologies of yeast proteins are shown. In each figure, full bars correspond to cocultures with *A. tumefaciens* 6N2 (pME6000) and dashed bars correspond to cocultures with *A. tumefaciens* 6N2 (pME6863). Y6N^QS+^_up_ and Y6N^QS-^_up_ subgroups are depicted in brown (A). Y6N^QS+^_dw_ and Y6N^QS-^_dw_ subgroups are depicted in blue (B).

## References

[1] J. Mansfield, S. Genin, S. Magori, V. Citovsky, M. Sriariyanum, P. Ronald, M. Dow, V. Verdier, S. V Beer, M.A. Machado, I. Toth, G. Salmond, G.D. Foster, Top 10 plant pathogenic bacteria in molecular plant pathology., Mol. Plant Pathol. 13 (2012) 614–29. https://doi.org/10.1111/j.1364-3703.2012.00804.x.

[2] F. Lassalle, T. Campillo, L. Vial, J. Baude, D. Costechareyre, D. Chapulliot, M. Shams, D. Abrouk, C. Lavire, C. Oger-Desfeux, F. Hommais, L. Guéguen, V. Daubin, D. Muller, X. Nesme, Genomic species are ecological species as revealed by comparative genomics in Agrobacterium tumefaciens, Genome Biol. Evol. 3 (2011) 762–781. https://doi.org/10.1093/gbe/evr070.

[3] Y. Dessaux, D. Faure, Quorum sensing and quorum quenching in Agrobacterium: A “Go/No Go system”?, Genes (Basel). 9 (2018) 210. https://doi.org/10.3390/genes9040210.

[4] C. Fuqua, S. Winans, E. Greenberg, Census and consensus in bacterial ecosystems: the LuxR-LuxI family of quorum-sensing transcriptional regulators, Annu. Rev. Microbiol. 50 (1996) 727–751.

[5] C. Fuqua, E.P. Greenberg, Listening in on bacteria: acyl-homoserine lactone signalling., Nat. Rev. Mol. Cell Biol. 3 (2002) 685–695. https://doi.org/10.1038/nrm907.

[6] I. Hwang, P.L. Li, L. Zhang, K.R. Piper, D.M. Cook, M.E. Tate, S.K. Farrand, TraI, a LuxI homologue, is responsible for production of conjugation factor, the Ti plasmid N-acylhomoserine lactone autoinducer., Proc. Natl. Acad. Sci. 91 (1994) 4639–4643. https://doi.org/10.1073/pnas.91.11.4639.

[7] K.M. Pappas, S.C. Winans, A LuxR-type regulator from Agrobacterium tumefaciens elevates Ti plasmid copy number by activating transcription of plasmid replication genes, Mol. Microbiol. 48 (2003) 1059–1073. https://doi.org/10.1046/j.1365-2958.2003.03488.x.

[8] Y.-X. Xing, C.-Y. Wei, Y. Mo, L.-T. Yang, S.-L. Huang, Y.-R. Li, Nitrogen-fixing and plant growth-promoting ability of two endophytic bacterial strains isolated from sugarcane stalks, Sugar Tech. 18 (2016) 373–379. https://doi.org/10.1007/s12355-015-0397-7.

[9] M. Fan, Z. Liu, L. Nan, E. Wang, W. Chen, Y. Lin, G. Wei, Isolation, characterization, and selection of heavy metal-resistant and plant growth-promoting endophytic bacteria from root nodules of Robinia pseudoacacia in a Pb/Zn mining area, Microbiol. Res. 217 (2018) 51–59. https://doi.org/10.1016/j.micres.2018.09.002.

[10] A.C.D. V Leguina, C. Nieto, H.F. Pajot, E. V Bertini, W. Mac Cormack, L.I. Castellanos de Figueroa, C.G. Nieto-Peñalver, Inactivation of bacterial quorum sensing signals N-acyl homoserine lactones is widespread in yeasts., Fungal Biol. 122 (2018) 52–62. https://doi.org/10.1016/j.funbio.2017.10.006.

[11] E. V. Bertini, Importancia de los mecanismos de quorum sensing en las interacciones entre microorganismos endofíticos. PhD Thesis. Universidad Nacional de Tucumán, 2018.

[12] E.V. Bertini, A.C. del V. Leguina, L.I. Castellanos de Figueroa, C.G. Nieto-Peñalver, Endophytic microorganisms Agrobacterium tumefaciens 6N2 and Meyerozyma guilliermondii 6N serve as models for the study of microbial interactions in colony biofilms, Rev. Argent. Microbiol. (2019). https://doi.org/10.1016/j.ram.2018.09.006.

[13] N. Mhedbi-Hajri, N. Yahiaoui, S. Mondy, N. Hue, F. Pélissier, D. Faure, Y. Dessaux, Transcriptome analysis revealed that a quorum sensing system regulates the transfer of the pAt megaplasmid in Agrobacterium tumefaciens, BMC Genomics. 17 (2016) 661. https://doi.org/10.1186/s12864-016-3007-5.

[14] C. Reimmann, N. Ginet, L. Michel, C. Keel, P. Michaux, V. Krishnapillai, M. Zala, K. Heurlier, D. Haas, K. Triandafillu, H. Harms, Genetically programmed autoinducer destruction reduces virulence gene expression and swarming motility in Pseudomonas aeruginosa PAO1, Microbiology. 148 (2002) 923–932.

[15] M. Cleene, The susceptibility of monocotyledons to Agrobacterium tumefaciens, J. Phytopathol. 113 (1985) 81–89. https://doi.org/10.1111/j.1439-0434.1985.tb00829.x.

[16] G. Suen, B.S. Goldman, R.D. Welch, Predicting prokaryotic ecological niches using genome sequence analysis., PLoS One. 2 (2007) e743. https://doi.org/10.1371/journal.pone.0000743.

[17] N. Calatrava-Morales, M. McIntosh, M.J. Soto, Regulation mediated by N-acyl homoserine lactone quorum sensing signals in the Rhizobium-legume symbiosis., Genes (Basel). 9 (2018) 263. https://doi.org/10.3390/genes9050263.

[18] G. Hao, T.J. Burr, Regulation of long-chain N-acyl-homoserine lactones in Agrobacterium vitis, J. Bacteriol. 188 (2006) 2173–83. https://doi.org/10.1128/JB.188.6.2173-2183.2006.

[19] S. Slater, J.C. Setubal, B. Goodner, K. Houmiel, J. Sun, R. Kaul, B.S. Goldman, S.K. Farrand, N. Almeida, T. Burr, E. Nester, D.M. Rhoads, R. Kadoi, T. Ostheimer, N. Pride, A. Sabo, E. Henry, E. Telepak, L. Cromes, A. Harkleroad, L. Oliphant, P. Pratt-Szegila, R. Welch, D. Wood, Reconciliation of sequence data and updated annotation of the genome of Agrobacterium tumefaciens C58, and distribution of a linear chromosome in the genus Agrobacterium, Appl. Environ. Microbiol. 79 (2013) 1414–1417. https://doi.org/10.1128/AEM.03192-12.

[20] D. Zheng, H. Zhang, S. Carle, G. Hao, M.R. Holden, T.J. Burr, A luxR homolog, aviR, in Agrobacterium vitis is associated with induction of necrosis on grape and a hypersensitive response on tobacco., Mol. Plant-Microbe Interact. 16 (2003) 650–8. https://doi.org/10.1094/MPMI.2003.16.7.650.

[21] S. Süle, L. Cursino, D. Zheng, H.C. Hoch, T.J. Burr, Surface motility and associated surfactant production in Agrobacterium vitis, Lett. Appl. Microbiol. 49 (2009) 596– 601. https://doi.org/10.1111/j.1472-765X.2009.02716.x.

[22] C.G. Nieto Penalver, F. Cantet, D. Morin, D. Haras, J.A. Vorholt, A plasmid-borne truncated luxI homolog controls quorum-sensing systems and extracellular carbohydrate production in Methylobacterium extorquens AM1, J. Bacteriol. 188 (2006) 7321–4. https://doi.org/10.1128/JB.00649-06.

[23] C. Wang, C. Yan, C. Fuqua, L.-H. Zhang, Identification and characterization of a second quorum-sensing system in Agrobacterium tumefaciens A6, J. Bacteriol. 196 (2014) 1403–1411. https://doi.org/10.1128/JB.01351-13.

[24] M.E. Wetzel, K.-S. Kim, M. Miller, G.J. Olsen, S.K. Farrand, Quorum-dependent mannopine-inducible conjugative transfer of an Agrobacterium opine-catabolic plasmid, J. Bacteriol. 196 (2014) 1031–1044. https://doi.org/10.1128/JB.01365-13.

[25] L. Leloup, E.-M. Lai, C. Kado, Identification of a chromosomal tra-like region in Agrobacterium tumefaciens, Mol. Genet. Genomics. 267 (2002) 115–123. https://doi.org/10.1007/s00438-002-0646-9.

[26] T. Netzker, J. Fischer, J. Weber, D.J. Mattern, C.C. König, V. Valiante, V. Schroeckh, A.A. Brakhage, Microbial communication leading to the activation of silent fungal secondary metabolite gene clusters., Front. Microbiol. 6 (2015) 299. https://doi.org/10.3389/fmicb.2015.00299.

[27] C. Grandclément, M. Tannières, S. Moréra, Y. Dessaux, D. Faure, Quorum quenching: role in nature and applied developments., FEMS Microbiol. Rev. 40 (2016) 86–116. https://doi.org/10.1093/femsre/fuv038.

[28] C.J. Wright, L.H. Burns, A.A. Jack, C.R. Back, L.C. Dutton, A.H. Nobbs, R.J. Lamont, H.F. Jenkinson, Microbial interactions in building of communities., Mol. Oral Microbiol. 28 (2013) 83–101. https://doi.org/10.1111/omi.12012.

[29] E. V Stabb, Could positive feedback enable bacterial pheromone signaling to coordinate behaviors in response to heterogeneous environmental cues?, MBio. 9 (2018). https://doi.org/10.1128/mBio.00098-18.

[30] C.B. Silveira, F.L. Rohwer, Piggyback-the-Winner in host-associated microbial communities., NPJ Biofilms Microbiomes. 2 (2016) 16010. https://doi.org/10.1038/npjbiofilms.2016.10.

[31] D. Tan, M.F. Hansen, L.N. de Carvalho, H.L. Røder, M. Burmølle, M. Middelboe, S. Lo Svenningsen, High cell densities favor lysogeny: induction of an H20 prophage is repressed by quorum sensing and enhances biofilm formation in Vibrio anguillarum., ISME J. 14 (2020) 1731–1742. https://doi.org/10.1038/s41396-020-0641-3.

[32] D. a Hogan, A. Vik, R. Kolter, A Pseudomonas aeruginosa quorum-sensing molecule influences Candida albicans morphology., Mol. Microbiol. 54 (2004) 1212–23. https://doi.org/10.1111/j.1365-2958.2004.04349.x.

[33] C. Boon, Y. Deng, L.-H. Wang, Y. He, J.-L. Xu, Y. Fan, S.Q. Pan, L.-H. Zhang, A novel DSF-like signal from Burkholderia cenocepacia interferes with Candida albicans morphological transition., ISME J. 2 (2008) 27–36. https://doi.org/10.1038/ismej.2007.76.

[34] B.M. Davis, R. Jensen, P. Williams, P. O’Shea, The interaction of N-acylhomoserine lactone quorum sensing signaling molecules with biological membranes: implications for inter-kingdom signaling., PLoS One. 5 (2010) e13522. https://doi.org/10.1371/journal.pone.0013522.

[35] F.E. Reyes-Turcu, K.H. Ventii, K.D. Wilkinson, Regulation and cellular roles of ubiquitin-specific deubiquitinating enzymes., Annu. Rev. Biochem. 78 (2009) 363– 397. https://doi.org/10.1146/annurev.biochem.78.082307.091526.

[36] H.G. Dohlman, S.L. Campbell, Regulation of large and small G proteins by ubiquitination., J. Biol. Chem. 294 (2019) 18613–18623. https://doi.org/10.1074/jbc.REV119.011068.

[37] S.C. Li, P.M. Kane, The yeast lysosome-like vacuole: endpoint and crossroads., Biochim. Biophys. Acta. 1793 (2009) 650–63. https://doi.org/10.1016/j.bbamcr.2008.08.003.

[38] M. Maurhofer, C. Reimmann, P. Schmidli-Sacherer, S. Heeb, D. Haas, G. Défago, Salicylic acid biosynthetic genes expressed in Pseudomonas fluorescens strain P3 improve the induction of systemic resistance in tobacco against Tobacco Necrosis Virus, Phytopathology. 88 (1998) 678–684. https://doi.org/10.1094/PHYTO.1998.88.7.678.

[39] P.D. Shaw, G. Ping, S.L. Daly, C. Cha, J.E. Cronan, K.L. Rinehart, S.K. Farrand, Detecting and characterizing N-acyl-homoserine lactone signal molecules by thinlayer chromatography., Proc. Natl. Acad. Sci. U. S. A. 94 (1997) 6036–41. https://doi.org/10.1073/pnas.94.12.6036.

[40] D. Vallenet, E. Belda, A. Calteau, S. Cruveiller, S. Engelen, A. Lajus, F. L. Fèvre, C. Longin, D. Mornico, D. Roche, Z. Rouy, G. Salvignol, C. Scarpelli, A.A. Thil Smith, M. Weiman, C. Médigue, MicroScope--an integrated microbial resource for the curation and comparative analysis of genomic and metabolic data., Nucleic Acids Res. 41 (2013) D636–D647. https://doi.org/10.1093/nar/gks1194.

[41] G.H. Van Domselaar, P. Stothard, S. Shrivastava, J.A. Cruz, A. Guo, X. Dong, P. Lu, D. Szafron, R. Greiner, D.S. Wishart, BASys: a web server for automated bacterial genome annotation., Nucleic Acids Res. 33 (2005) W455–W459. https://doi.org/10.1093/nar/gki593.

[42] D.H. Figurski, D.R. Helinski, Replication of an origin-containing derivative of plasmid RK2 dependent on a plasmid function provided in trans., Proc. Natl. Acad. Sci. U. S. A. 76 (1979) 1648–1652. https://doi.org/10.1073/pnas.76.4.1648.

[43] C. Cha, P. Gao, Y.C. Chen, P.D. Shaw, S.K. Farrand, Production of acyl-homoserine lactone quorum-sensing signals by gram-negative plant-associated bacteria., Mol. Plant-Microbe Interact. 11 (1998) 1119–1129. https://doi.org/10.1094/MPMI.1998.11.11.1119.

[44] L. Ravn, A.B. Christensen, S. Molin, M. Givskov, L. Gram, Methods for detecting acylated homoserine lactones produced by Gram-negative bacteria and their application in studies of AHL-production kinetics, J. Microbiol. Methods. 44 (2001) 239–251. https://doi.org/10.1016/S0167-7012(01)00217-2.

[45] Y. Perez-Riverol, A. Csordas, J. Bai, M. Bernal-Llinares, S. Hewapathirana, D.J. Kundu, A. Inuganti, J. Griss, G. Mayer, M. Eisenacher, E. Pérez, J. Uszkoreit, J. Pfeuffer, T. Sachsenberg, Ş. Yılmaz, S. Tiwary, J. Cox, E. Audain, M. Walzer, A.F. Jarnuczak, T. Ternent, A. Brazma, J.A. Vizcaíno, The PRIDE database and related tools and resources in 2019: improving support for quantification data, Nucleic Acids Res. 47 (2019) D442–D450. https://doi.org/10.1093/nar/gky1106.

