## Supplementary material for "Insight in the quorum sensing-driven lifestyle of the non-pathogenic *Agrobacterium tumefaciens* 6N2 and the interactions with the yeast *Meyerozyma guilliermondii*": Supp. Materials

**Genomic sequencing, *de novo* assembly and annotation.** Genome sequence was obtained utilizing single-molecule real-time sequencing technology (Pacific Biosciences). SMRTbell template libraries were prepared with Template Preparation Kit (Pacific Biosciences), annealed with Template Binding Kit (Pacific Biosciences) and bound to P6 DNA polymerase. For enhanced efficiency, bound complexes were immobilized into Magbeads (Pacific Biosciences). Sequence collection was carried out in SMRT cells using P6/C4 chemistry. Reads less than 500 bp were filtered off and minimum polymerase read quality was set at 0.75. Reads assembled using RS_HGAP_Assembly 3.0. Genome annotation was performed with the MicroScope platform [1], BASys [2] and RAST [3]. Genome visualization was performed with CGViewer [4]. Cumulative GC skew was analyzed with GenSkew (http://genskew.csb.univie.ac.at/). Prophages were identified with PHASTER [5], type III, IV and VI secretion systems with T346Hunter [6], genomic islands with IslandViewer 4 [7] and integrative and conjugative elements (ICEs) with ICEfinder [8]. Genome sequences were deposited in Genbank under accession numbers CP072308 and CP072309.

**Identification and characterization of QS genes.** BLAST searches were performed on strain 6N2 genome utilizing as query the *traI* and *traR* of *A. fabrum* strain C58 and related microorganisms. Protein domain architecture was determined with Pfam [9] and SMART [10]. Topology of QS systems was compared to those of *Agrobacterium* spp., with complete genomes available at the Joint Genome Institute (JGI) database. Alignments of LuxI and LuxR orthologues were obtained with MAFFT [11], and edited when required with MEGA [12]. Identity matrices were constructed with BioEdit [13] and visualized in heatmaps with MORPHEUS (https://software.broadinstitute.org/morpheus). Synteny around the QS systems was visualized with SimpleSinteny [14].

**Preparation of protein extracts.** *A. tumefaciens* 6N2 (pME6000), *A. tumefaciens* 6N2 (pME6863), and *M. guilliermondii* 6N were precultured aerobically overnight in NB broth at 30 ºC. Precultures were centrifuged at 7,000 *g*, supernatants were discarded and cell pellets were resuspended in sterile NB broth. Suspensions were utilized to prepare pure and dual cultures of both microorganisms. Two hundred and fifty mL flasks containing 20 mL of NB broth were inoculated at an initial concentration of ~10^7^ CFU mL^-1^ for *A. tumefaciens* 6N2 (pME6000) or *A. tumefaciens* 6N2 (pME6863), and ~10^6^ CFU mL^-1^ for *M. guilliermondii* 6N. Dual cultures of *A. tumefaciens* 6N2 (pME6000) plus the yeast, and *A. tumefaciens* 6N2 (pME6863) plus the yeast were prepared with those cell densities. Flasks were incubated aerobically at 30 ºC for 24 h until late exponential growth phase, then centrifuged at 10,000 *g* for 10 min and washed twice with PBS buffer. Protein extracts were obtained with YPX extraction kit (EXPEDEON). Protein concentrations were determined with the QuantiPro BCA (SigmaAldrich). Three independent samples were analyzed for each pure or mixed culture.

**Proteomics acquisitions.**

Protein samples were trypsin digested with FASP (Protein Digestion Kit, EXPEDEON) following manufacture instructions. Digested peptide mixtures were analyzed by a Q-Exactive mass spectrometer coupled to an Easy-nLC system (both from Thermo Scientific). Chromatographic separations were performed with the following parameters: Acclaim PepMap100 C18 precolumn (2 cm, 75 µm i.d., 3 µm, 100 Å), Pepmap-RSLC Proxeon C18 column (50 cm, 75 µm i.d., 2 µm, 100 Å), 300 nL min^-1^ flow, gradient rising from 95% solvent A (water, 0.1% formic acid) to 35% solvent B (100% acetonitrile, 0.1% formic acid) in 98 min followed by a column regeneration for 23 min giving a total run time of 2 hours. Peptides were analyzed in the Orbitrap in full ion scan mode at a resolution of 70,000 (at *m*/*z* 200) with a mass range of *m*/*z* 375–1500. Fragments were obtained by Higher-energy Collisional Dissociation (HCD) activation with a collisional energy of 28 %, an isolation width of 1.4 Da. MS/MS data were acquired in the Orbitrap cell in a data-dependent mode in which the 20 most intense precursor ions were fragmented, with a dynamic exclusion of 20 seconds and a resolution of 17,500. The maximum ion accumulation times were set to 50 ms for MS acquisition and 45 ms for MS/MS acquisition.

**Proteomics data and bioinformatic analysis.** All MS/MS data were processed with Proteome Discoverer 2.1 (Thermo Scientific) coupled to an in-house Mascot search server (Matrix Science, Boston, MA; version 2.5.1). The mass tolerance was set to 6 ppm for precursor ions and 0.02 Da for fragments. The following modifications were used in variable modifications: oxidation (M), phosphorylation (STY), acetylation (N-term), and carbamidomethylation (C) in fixed modification. The maximum number of missed cleavages by trypsin was limited to two. MS/MS data were analyzed with an *ad hoc* database composed of proteins obtained from the annotation of the *A. tumefaciens* strain 6N2 genome, and 5923 proteins from *M. guilliermondii* ATCC 6260 obtained from UniProt. Peptide identifications were validated by using a False Discovery Rate (FDR) threshold of 0.01 calculated with the Percolator algorithm. Peptides and proteins abundance variations were measured by using Progenesis QI for proteomics software (version 4.0, Waters) and a hi-3 method for which the 3-most-abundant peptides were considered for each protein quantification. Proteins showing a fold change (FC) ≥ 1.5 and an ANOVA p≤0.05 were considered as differentially accumulated. For mixed culture, quantification of *A. tumefaciens* and *M. guilliermondii* proteins were considered only after an abundance normalization based on total ion current (TIC) and with exclusively *A. tumefaciens* and *M. guilliermondii* proteins respectively. Proteins showing a fold change (FC) ≥ 1.5 and an ANOVA p≤0.05 were considered as differentially accumulated. Proteins were functionally categorized after mapping in eggNOG [15]. Bacterial proteins were blasted against the UniProt database and corresponding GO annotations of the best hits were retrieved. For fungal proteins, GO annotations were directly obtained from UniProt [16]. Redundant GO terms were eliminated with CateGOrizer [17].

Complete datasets are available in the ProteomeXchange Consortium via the PRIDE [18] partner repository with the identifier PXD025730. Mass spectrometer output files are available as .raw files; protein identification files are available as .msf files; PEAK list files are available as .mgf files; quantification files are available as .xlsx files.

**References**

[1]	D. Vallenet, E. Belda, A. Calteau, S. Cruveiller, S. Engelen, A. Lajus, F. Le Fèvre, C. Longin, D. Mornico, D. Roche, Z. Rouy, G. Salvignol, C. Scarpelli, A.A. Thil Smith, M. Weiman, C. Médigue, MicroScope--an integrated microbial resource for the curation and comparative analysis of genomic and metabolic data., Nucleic Acids Res. 41 (2013) D636–D647. https://doi.org/10.1093/nar/gks1194.

[2]	G.H. Van Domselaar, P. Stothard, S. Shrivastava, J.A. Cruz, A. Guo, X. Dong, P. Lu, D. Szafron, R. Greiner, D.S. Wishart, BASys: a web server for automated bacterial genome annotation., Nucleic Acids Res. 33 (2005) W455–W459. https://doi.org/10.1093/nar/gki593.

[3]	R. Overbeek, R. Olson, G.D. Pusch, G.J. Olsen, J.J. Davis, T. Disz, R.A. Edwards, S. Gerdes, B. Parrello, M. Shukla, V. Vonstein, A.R. Wattam, F. Xia, R. Stevens, The SEED and the Rapid Annotation of microbial genomes using Subsystems Technology (RAST)., Nucleic Acids Res. 42 (2014) D206–D214. https://doi.org/10.1093/nar/gkt1226.

[4]	J.R. Grant, P. Stothard, The CGView Server: a comparative genomics tool for circular genomes, Nucleic Acids Res. 36 (2008) W181–W184. https://doi.org/10.1093/nar/gkn179.

[5]	D. Arndt, J.R. Grant, A. Marcu, T. Sajed, A. Pon, Y. Liang, D.S. Wishart, PHASTER: a better, faster version of the PHAST phage search tool., Nucleic Acids Res. 44 (2016) W16–W21. https://doi.org/10.1093/nar/gkw387.

[6]	P.M. Martínez-García, C. Ramos, P. Rodríguez-Palenzuela, T346Hunter: a novel web-based tool for the prediction of type III, type IV and type VI secretion systems in bacterial genomes., PLoS One. 10 (2015) e0119317. https://doi.org/10.1371/journal.pone.0119317.

[7]	C. Bertelli, M.R. Laird, K.P. Williams, B.Y. Lau, G. Hoad, G.L. Winsor, F.S. Brinkman, IslandViewer 4: expanded prediction of genomic islands for larger-scale datasets, Nucleic Acids Res. 45 (2017) W30–W35. https://doi.org/10.1093/nar/gkx343.

[8]	M. Liu, X. Li, Y. Xie, D. Bi, J. Sun, J. Li, C. Tai, Z. Deng, H.-Y. Ou, ICEberg 2.0: an updated database of bacterial integrative and conjugative elements., Nucleic Acids Res. 47 (2019) D660–D665. https://doi.org/10.1093/nar/gky1123.

[9]	S. El-Gebali, J. Mistry, A. Bateman, S.R. Eddy, A. Luciani, S.C. Potter, M. Qureshi, L.J. Richardson, G.A. Salazar, A. Smart, E.L.L. Sonnhammer, L. Hirsh, L. Paladin, D. Piovesan, S.C.E. Tosatto, R.D. Finn, The Pfam protein families database in 2019, Nucleic Acids Res. 47 (2019) D427–D432. https://doi.org/10.1093/nar/gky995.

[10]	I. Letunic, P. Bork, 20 years of the SMART protein domain annotation resource, Nucleic Acids Res. 46 (2018) D493–D496. https://doi.org/10.1093/nar/gkx922.

[11]	K. Katoh, D.M. Standley, MAFFT multiple sequence alignment software version 7: improvements in performance and usability, Mol. Biol. Evol. 30 (2013) 772–780. https://doi.org/10.1093/molbev/mst010.

[12]	S. Kumar, G. Stecher, M. Li, C. Knyaz, K. Tamura, MEGA X: molecular evolutionary genetics analysis across computing platforms., Mol. Biol. Evol. 35 (2018) 1547–1549. https://doi.org/10.1093/molbev/msy096.

[13]	T.A. Hall, BioEdit: a user-friendly biological sequence alignment editor and analysis program for Windows 95/98/NT, Nucl. Acids Symp. Ser. 41 (1999) 95–98.

[14]	D. Veltri, M.M. Wight, J.A. Crouch, SimpleSynteny: a web-based tool for visualization of microsynteny across multiple species., Nucleic Acids Res. 44 (2016) W41–W45. https://doi.org/10.1093/nar/gkw330.

[15]	J. Huerta-Cepas, D. Szklarczyk, D. Heller, A. Hernández-Plaza, S.K. Forslund, H. Cook, D.R. Mende, I. Letunic, T. Rattei, L.J. Jensen, C. von Mering, P. Bork, eggNOG 5.0: a hierarchical, functionally and phylogenetically annotated orthology resource based on 5090 organisms and 2502 viruses, Nucleic Acids Res. 47 (2019) D309–D314. https://doi.org/10.1093/nar/gky1085.

[16]	UniProt Consortium, Update on activities at the Universal Protein Resource (UniProt) in 2013., Nucleic Acids Res. 41 (2013) D43–D47. https://doi.org/10.1093/nar/gks1068.

[17]	H. Zhi-Liang, J. Bao, J. Reecy, A web-based program to batch analyze gene ontology classification categories, Online J. Bioinforma. 9 (2008) 108–112.

[18]	Y. Perez-Riverol, A. Csordas, J. Bai, M. Bernal-Llinares, S. Hewapathirana, D.J. Kundu, A. Inuganti, J. Griss, G. Mayer, M. Eisenacher, E. Pérez, J. Uszkoreit, J. Pfeuffer, T. Sachsenberg, Ş. Yılmaz, S. Tiwary, J. Cox, E. Audain, M. Walzer, A.F. Jarnuczak, T. Ternent, A. Brazma, J.A. Vizcaíno, The PRIDE database and related tools and resources in 2019: improving support for quantification data, Nucleic Acids Res. 47 (2019) D442–D450. https://doi.org/10.1093/nar/gky1106.
