## Supplementary figures and images for "Insight in the quorum sensing-driven lifestyle of the non-pathogenic *Agrobacterium tumefaciens* 6N2 and the interactions with the yeast *Meyerozyma guilliermondii*"

### Suppl. Fig. 1

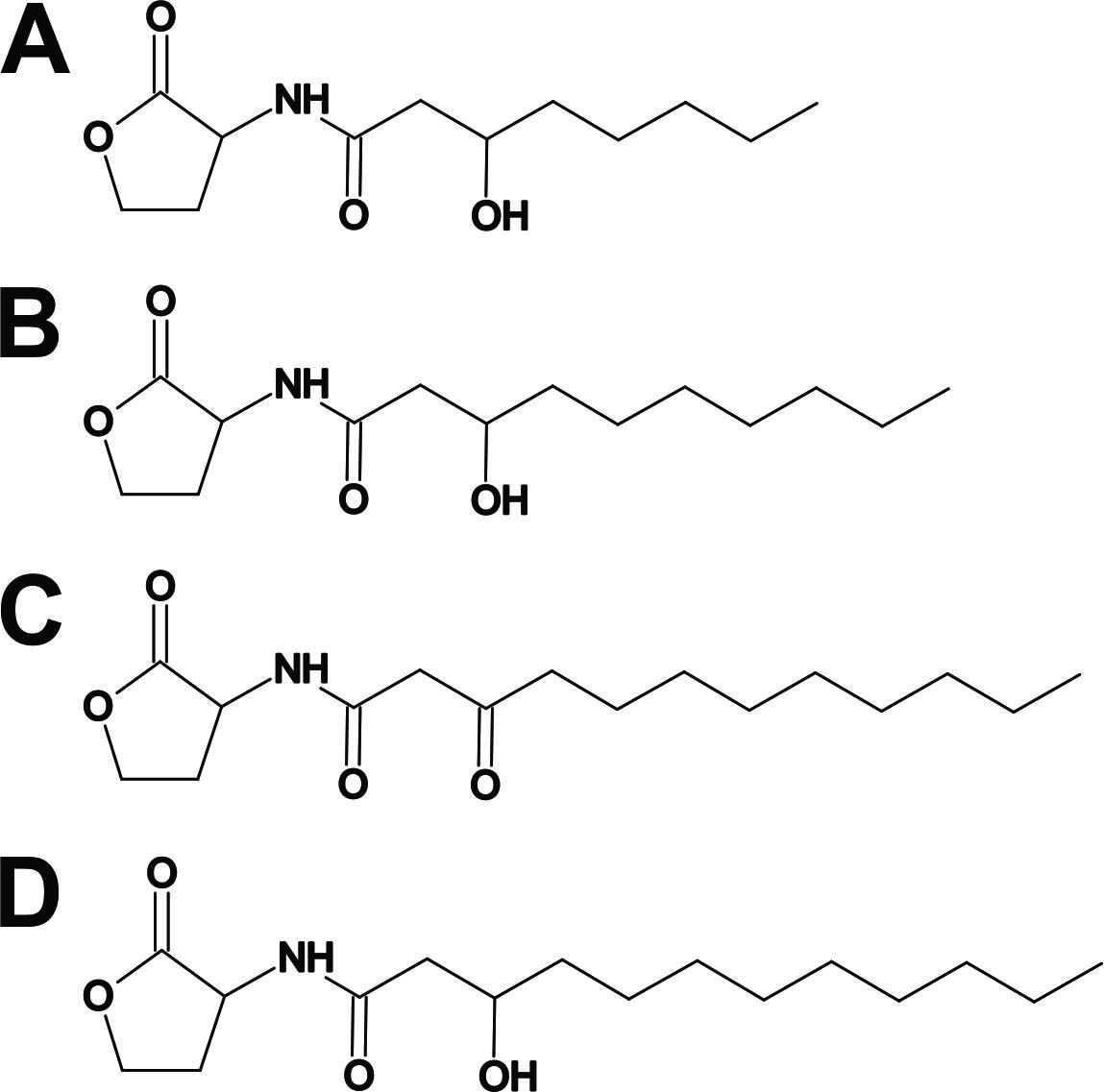

### Suppl. Fig. 2

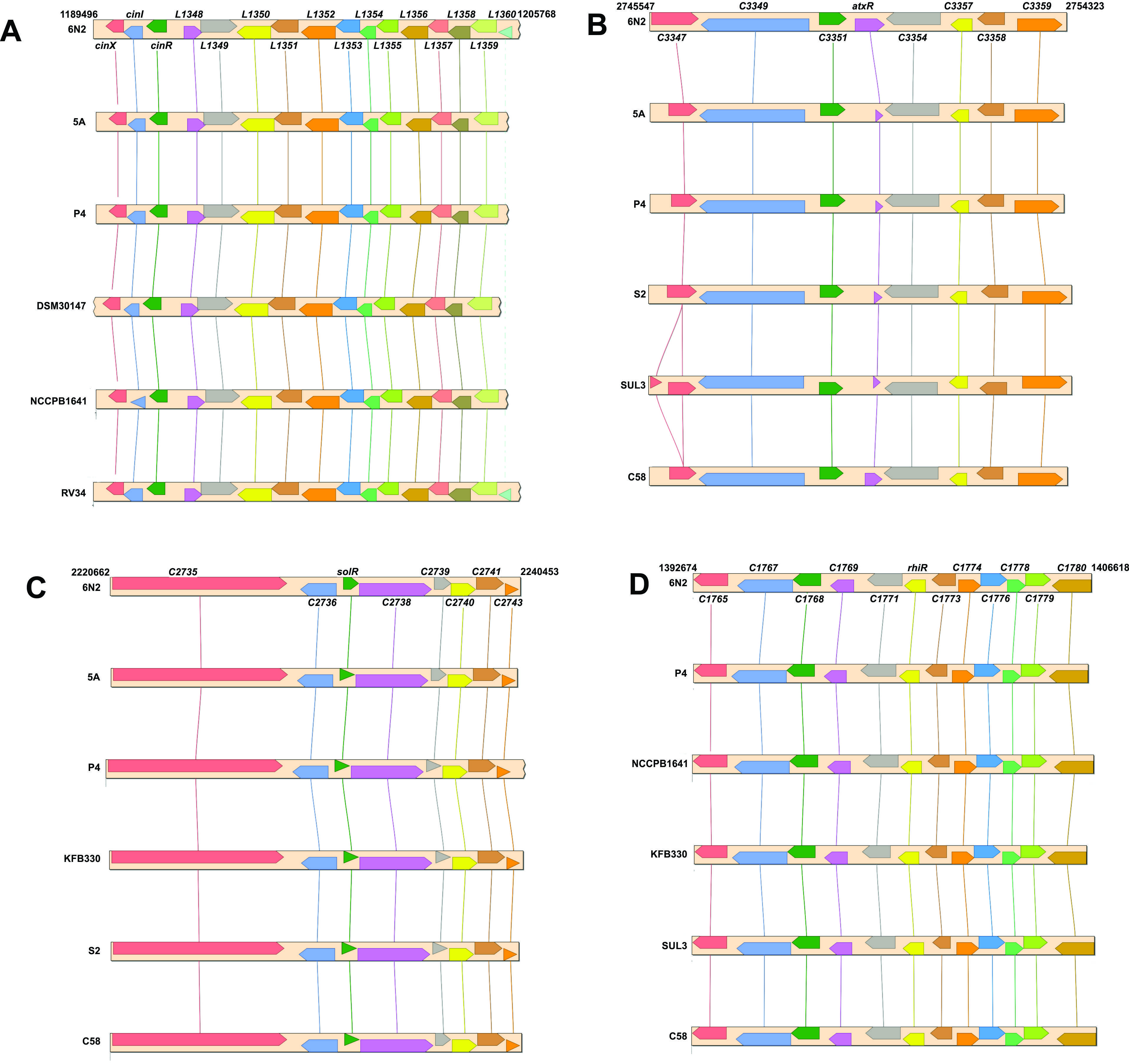

### Suppl. Fig. 3A

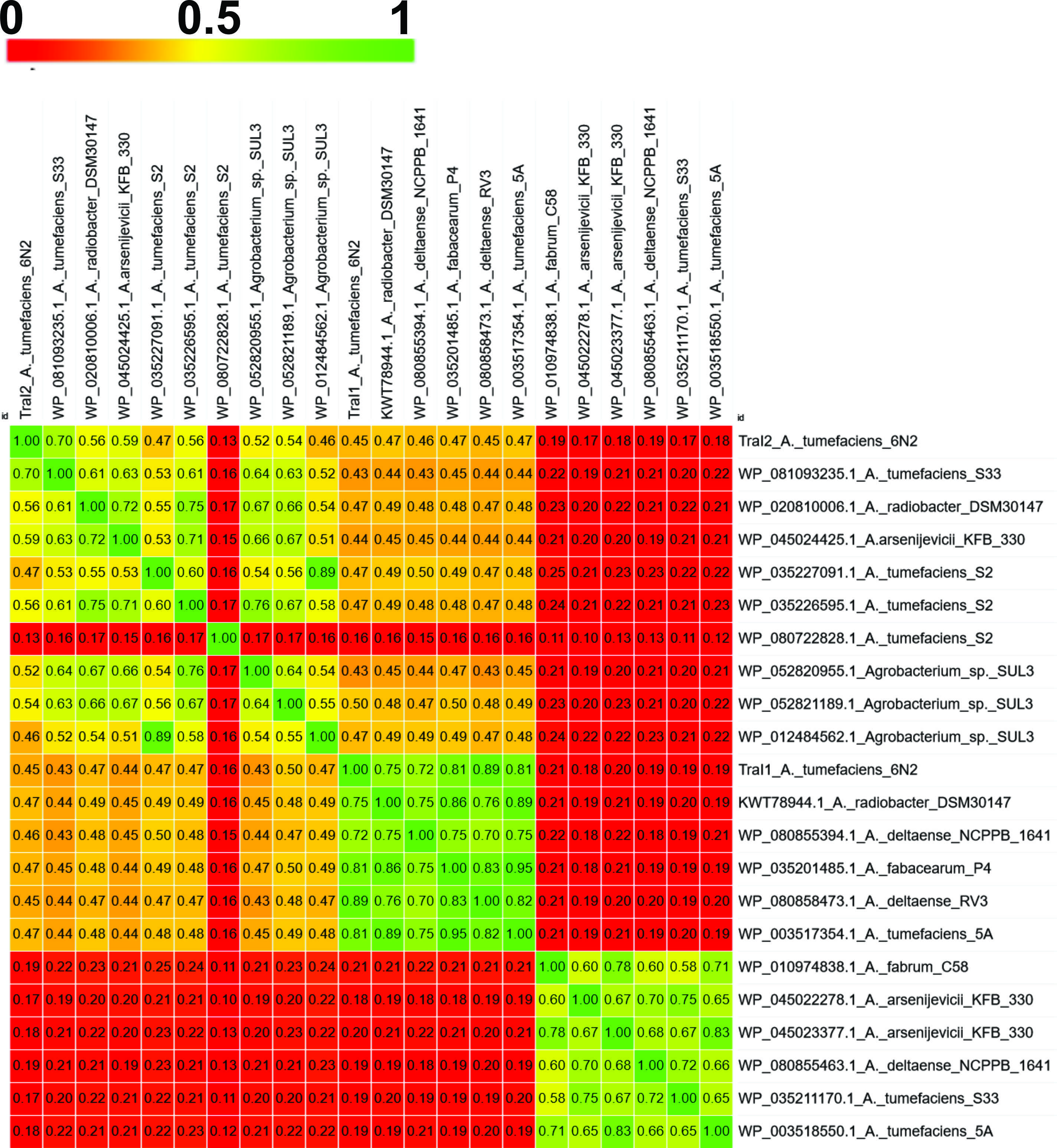

### Suppl. Fig. 3B

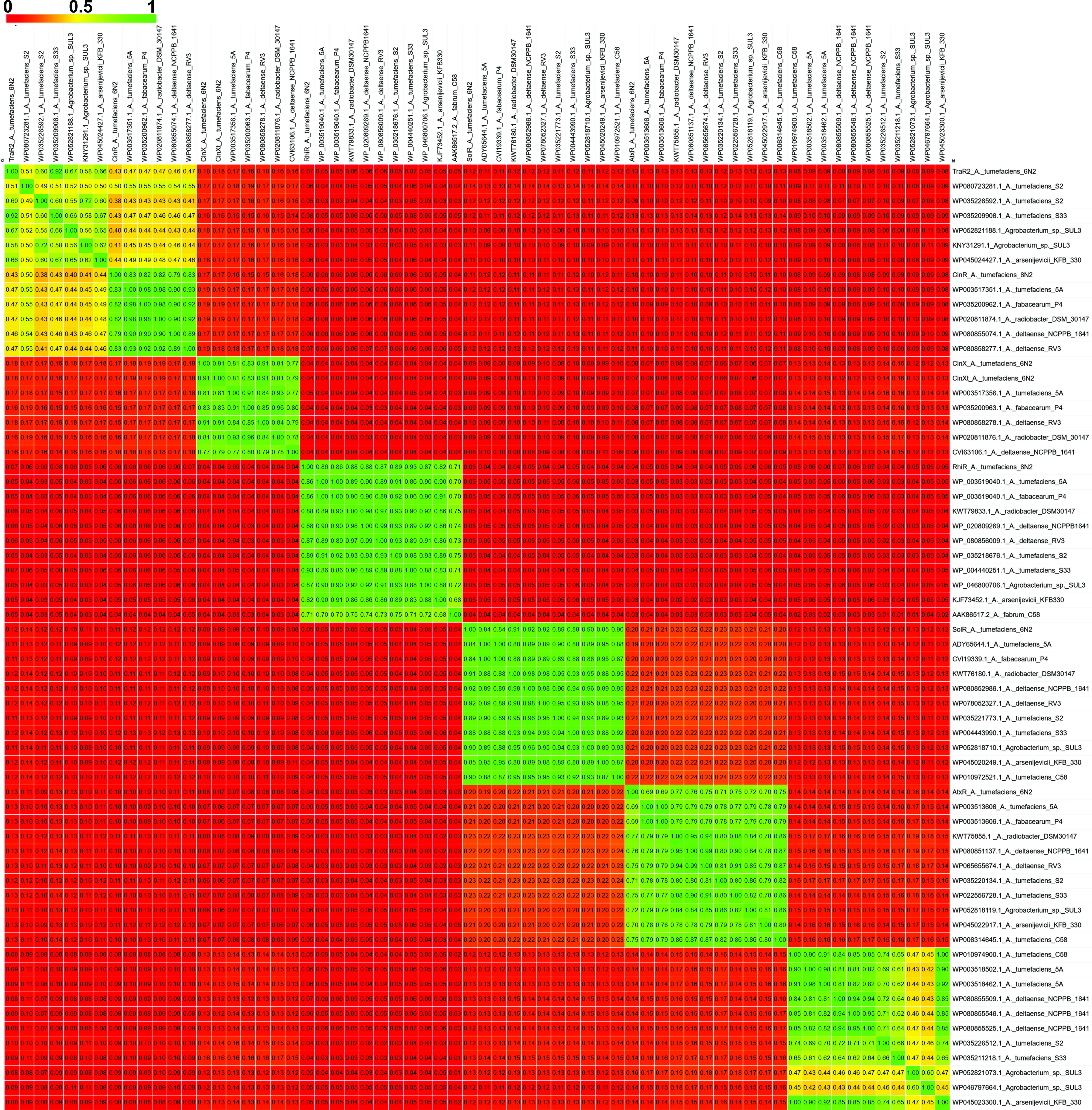

### Suppl. Fig. 4

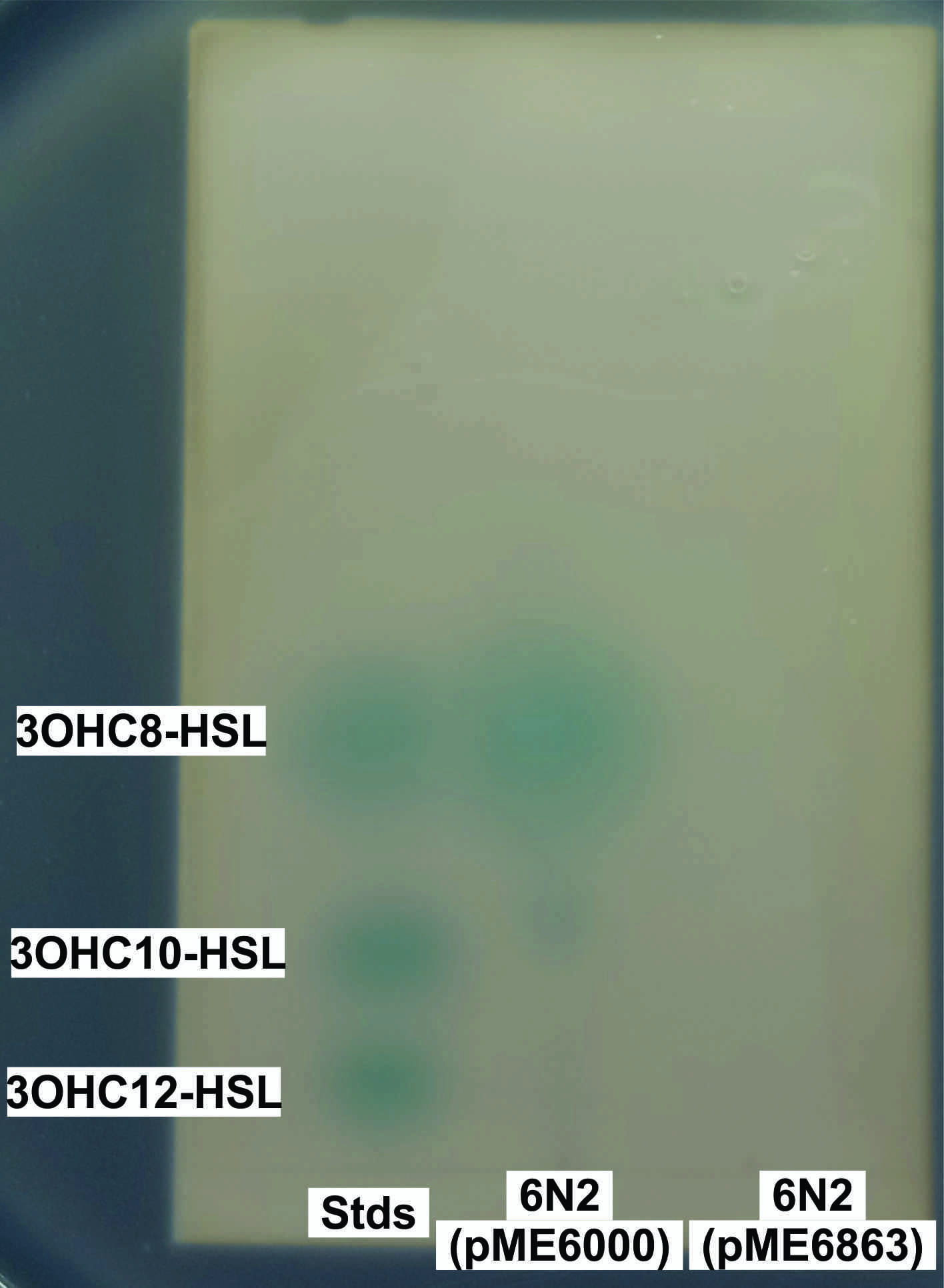

### Suppl. Fig. 5

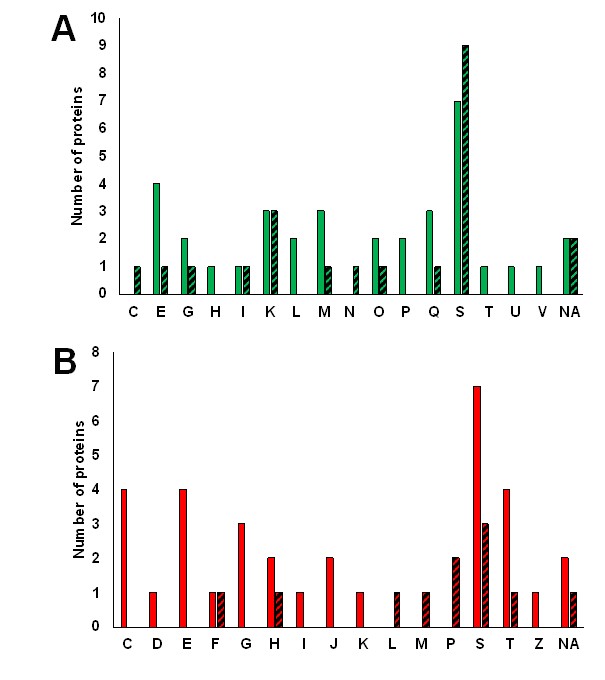

### Suppl. Fig. 6

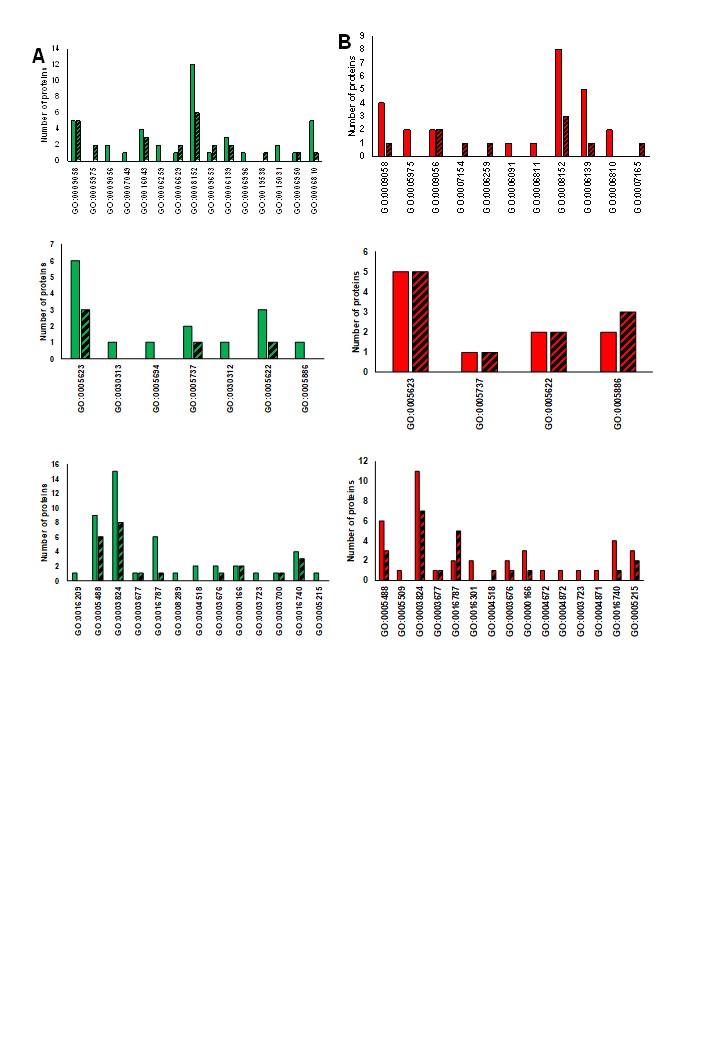

### Suppl. Fig. 7

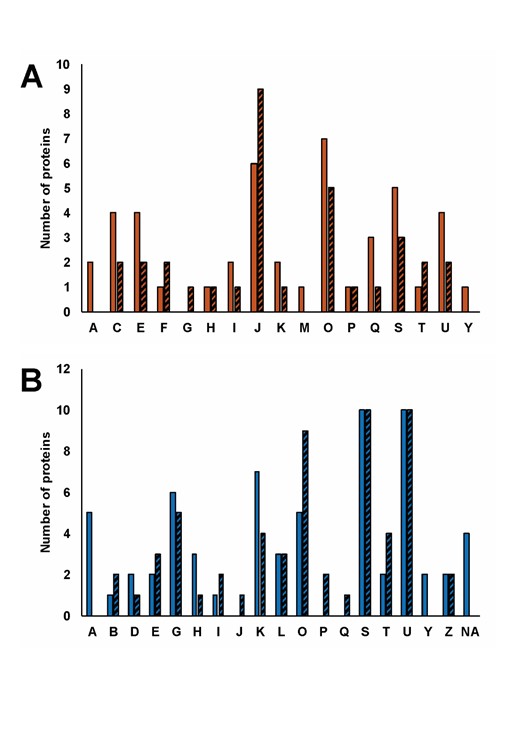

### Suppl. Fig. 8

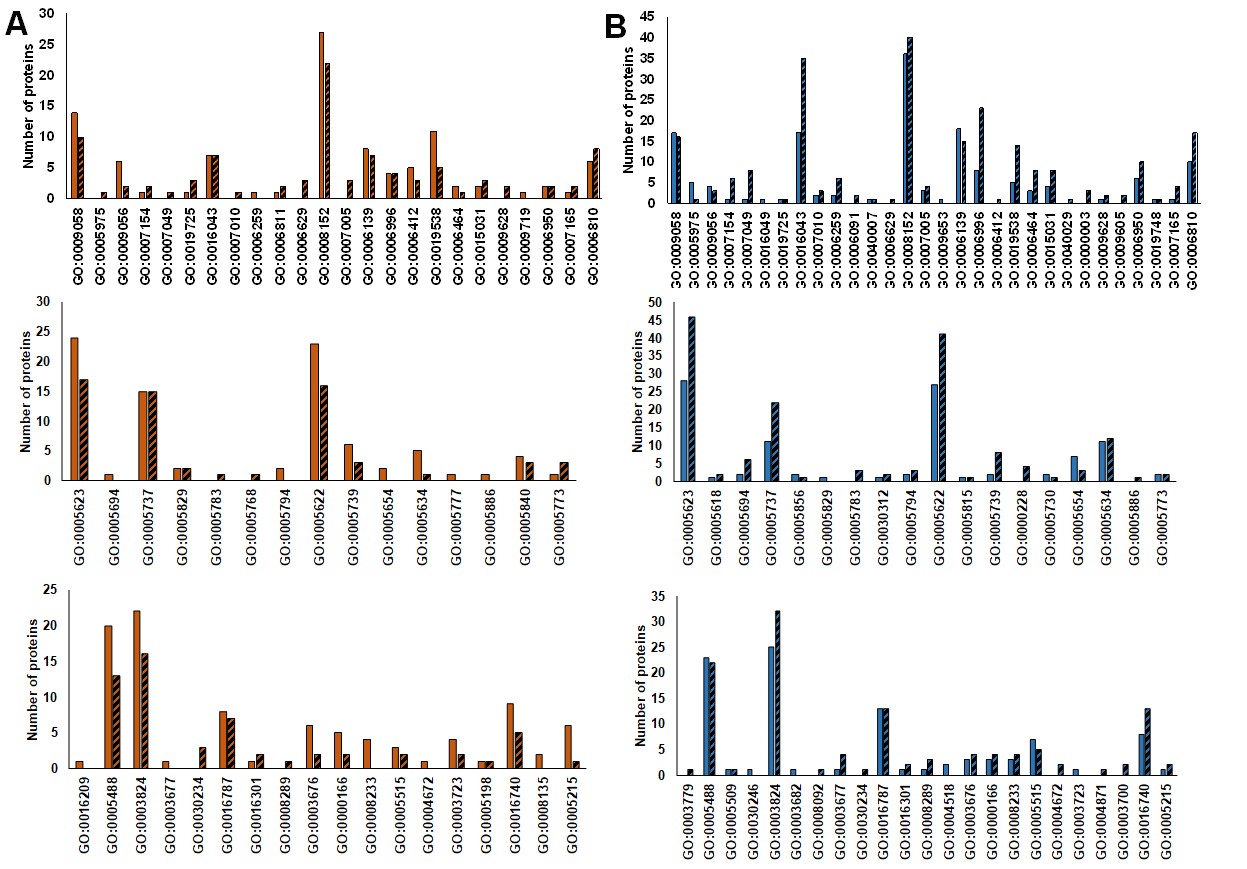
